# Mouse Nuclear RNAi-defective 2 Promotes Splicing of Weak 5’ Splice Sites

**DOI:** 10.1101/2022.01.25.477700

**Authors:** Matyas Flemr, Michaela Schwaiger, Daniel Hess, Vytautas Iesmantavicius, Alex Charles Tuck, Fabio Mohn, Marc Bühler

**Affiliations:** Friedrich Miescher Institute for Biomedical Research, Maulbeerstrasse 66, 4058 Basel, Switzerland; Swiss Institute of Bioinformatics, 4058 Basel, Switzerland; University of Basel, Petersplatz 10, 4003 Basel, Switzerland

**Keywords:** NRDE2, CCDC174, CLIP, RNA binding, U1 snRNP, weak 5’splice site, splice site selection

## Abstract

Removal of introns during pre-mRNA splicing, which is central to gene expression, initiates by base pairing of U1 snRNA with a 5’ splice site (5’SS). In mammals, many introns contain weak 5’SSs that are not efficiently recognized by the canonical U1 snRNP, suggesting alternative mechanisms exist. Here, we develop a cross-linking immunoprecipitation coupled to a high-throughput sequencing method, BCLIP-seq, to identify NRDE2 (Nuclear RNAi defective-2) and CCDC174 (Coiled-Coil Domain-Containing 174) as novel RNA-binding proteins in mouse ES cells that associate with U1 snRNA and unspliced 5’SSs. Both proteins bind directly to U1 snRNA independently of canonical U1 snRNP specific proteins, and they are required for the selection and effective processing of weak 5’SSs. Our results reveal that mammalian cells use non-canonical splicing factors bound directly to U1 snRNA to effectively select suboptimal 5’SS sequences in hundreds of genes, promoting proper splice site choice and accurate pre-mRNA splicing.

## INTRODUCTION

Accurate removal of noncoding intronic sequences during pre-mRNA splicing is a prerequisite for eukaryotic gene expression. Most introns are excised by the major spliceosome, a dynamic macromolecular assembly consisting of five small nuclear ribonucleoprotein particles (U1, U2, U4, U5, and U6 snRNPs) and a myriad of associated splicing factors (Wahl et al., 2009). The highly structured snRNA components of snRNPs bind complementary sequences in pre-mRNA that define and position the splice sites to form a catalytic spliceosome, in which two consecutive transesterification reactions liberate the intron and allow two neighboring exons to be ligated (Kastner et al., 2019; Shi, 2017; Wilkinson et al., 2019).

5’ splice site (5’SS) recognition by the U1 snRNP represents the first step of spliceosome assembly. 5’SSs comprise a short motif that is recognized by a complementary sequence at the 5’ end of the U1 snRNA (Kondo et al., 2015; Plaschka et al., 2018). However, major differences exist between model organisms in which pre-mRNA splicing has been investigated. In budding yeast, an almost invariant 5’SS motif is bound by a U1 snRNP containing a long U1 snRNA that triggers spliceosome assembly and rapid co-transcriptional splicing shortly after the intron has been transcribed (Carrillo Oesterreich et al., 2016; Lacadie and Rosbash, 2005). In contrast, other eukaryotes, ranging from fission yeast to mammals, encode much shorter U1 snRNAs packaged in smaller snRNPs that must cope with a highly degenerate 5’SS motif (Fair and Pleiss, 2017). The sequence variability of 5’SSs can decrease U1 binding affinity, resulting in weak 5’SSs with reduced splicing efficiency (Roca et al., 2013). Most mammalian genes contain multiple introns with several potential splice sites, among which the weak 5’SSs are often associated with alternative splicing patterns (Boutz et al., 2015; Drexler et al., 2020; Lee and Rio, 2015). These include alternative exon inclusion, exon skipping and intron retention, which can all possess regulatory functions by generating different protein isoforms or ensuring timely expression of mature transcripts (Mauger et al., 2016; Naro et al., 2017; Ule and Blencowe, 2019). However, alternative processing of weak 5’SSs can also lead to nonfunctional transcripts targeted for degradation by RNA surveillance pathways (Bresson et al., 2015; Davidson et al., 2012; Peck et al., 2019). Proper choice and efficient splicing of weak 5’SSs are therefore crucial for accurate gene expression, i.e. mutations decreasing splice site strength or impairing activity of splicing factors are a frequent cause of human genetic disorders (Anna and Monika, 2018; Scotti and Swanson, 2016; Wickramasinghe et al., 2015).

Due to the limited complementarity to U1 snRNA, weak 5’SSs are not efficiently recognized by the canonical U1 snRNP consisting of the U1 snRNA, a set of seven Sm proteins forming a ring structure common to other U snRNPs, and three U1-specific proteins U1A, U1C and U1-70K (Kondo et al., 2015). Therefore, additional splicing factors must be involved in efficient U1 binding and processing of weak 5’SSs. Despite a plethora of known splicing factors, additional proteins with uncharacterized splicing-related functions continue to be discovered. For example, NRDE2 (Nuclear RNAi defective-2) is a conserved protein with homologs detected in eukaryotes ranging from fission yeast to human, but absent in budding yeast. The fission yeast homolog Nrl1 has been shown to form a complex with Mtl1 (MTREX in mammals) and Ctr1 proteins (CCDC174 in mammals, for Coiled-Coil Domain-Containing 174) and interact with core splicing factors. The Nrl1-Ctr1-Mtl1 complex has been proposed to regulate splicing of cryptic introns and target unspliced transcripts for degradation by the nuclear exosome (Lee et al., 2013; Zhou et al., 2015). In contrast, mammalian NRDE2 has been suggested to reduce exosome activity by inhibiting the MTREX helicase (Wang et al., 2019). A NRDE2 interaction with CCDC174 has thus far not been detected (Richard et al., 2018). Nevertheless, human CCDC174 was shown to interact with a splicing-related exon junction complex (EJC) component EIF4A3 (Volodarsky et al., 2015). NRDE2 was also found to interact with splicing factors and its depletion led to the retention of weakly spliced introns (Jiao et al., 2019). However, potential mechanisms by which NRDE2 and CCDC174 might contribute to pre-mRNA splicing remain unknown, partly due to a lack of any predicted functional domains in these two largely uncharacterized proteins.

Here we combined genome engineering in mouse embryonic stem cells (mESCs) with proteomics and genomics approaches to analyze the function of NRDE2 in mammalian cells. In order to reliably detect potential RNA substrates of NRDE2 with high sensitivity and at a transcriptome-wide level, we have developed a modified cross-linking immunoprecipitation coupled to next-generation sequencing method. This revealed that mouse NRDE2 together with CCDC174 bind to U1 snRNAs, which target NRDE2 to 5’SSs. At weak 5’SSs with suboptimal U1 complementarity, NRDE2 and CCDC174 are necessary for correct splice site choice and efficient pre-mRNA splicing. Thus, we reveal that mammalian cells use an alternative strategy involving direct U1 snRNA interaction with non-canonical splicing factors to select and splice introns with suboptimal 5’SSs.

## RESULTS

### NRDE2 Is Required for Cell Growth Independently of Its Protein Interactors

Using our previously established mESC line expressing 4-hydroxytamoxifen (4OHT) inducible Cre recombinase and bacterial BirA ligase (Flemr and Bühler, 2015), we generated inducible *Nrde2* knockout cells. These cells were further edited to introduce homozygous *Nrde2* mutations and to tag NRDE2 with a fluorescent protein for live cell imaging or with a composite 3xFLAG-AviTag (3A tag) for tandem FLAG-Streptavidin purifications. The knockout and tagging approach was used on other endogenous genes in this study (Figure S1A and Table S1), which also included fusions with the 2xHA-FKBP12^F36V^ domain (dTAG) for rapid protein depletion by the dTAG-13 compound (Nabet et al., 2018). We assessed potential consequences of the tagging approach on protein function by comparing RNA-seq gene expression profiles with that of untagged cells, which remained unchanged for the proteins we report hereinafter.

In NRDE2 loss-of-function experiments the 4OHT-induced knockout of *Nrde2* resulted in growth arrest (Figure S1B), revealing NRDE2 has an essential function in cellular growth. To understand the basis for this phenotype, we mapped the NRDE2 protein-protein interaction (PPI) network by performing tandem affinity purifications of 3A-NRDE2 under mild native conditions coupled with mass spectrometry (nTAP-MS). The NRDE2 interactome was dominated by ribosomal proteins, splicing factors, and included the complete nuclear exosome (Figure 1A). The nuclear exosome-associated RNA helicase MTREX (formerly MTR4) was the highest scoring interactor, consistent with published data (Wang et al., 2019). Reciprocal co-immunoprecipitation (co-IP) of 3A-MTREX confirmed this interaction and NRDE2 truncation experiments revealed MTREX interacts with NRDE2 within a region including amino acids 101-200 (Figure S1C). Four conserved aspartates within this region (Figure S1D) resembled an MTREX Arch interaction motif (Thoms et al., 2015) and a single D174R point mutation in NRDE2 substantially reduced the interaction with MTREX (Figure S1E). NRDE2-D174R protein levels were lower compared to wild-type NRDE2 (Figure S1F), suggesting that MTREX controls NRDE2 stability. This was further supported by reduced NRDE2 levels upon *Mtrex* knockout (Figure S1G), which is in line with previously published data from human cells (Wang et al., 2019). However, the N-terminal 200 amino-acid truncation (NRDE2Δ200) was expressed at a comparable level to wild-type NRDE2. Thus, the N-terminal region is required for NRDE2 destabilization in the absence of MTREX. MTREX and nuclear exosome components were lost from the NRDE2-D174R nTAP-MS PPI network, whereas ribosomal proteins and several splicing factors were still present (Figure S1H). Strikingly, we did not detect any PPIs for NRDE2Δ200 under these conditions (Figure 1B), indicating the N-terminal region of NRDE2 is essential for interactions with other proteins. The expression of NRDE2-D174R or NRDE2Δ200 promoted cell viability, albeit at a slower growth rate in the latter case, with cell growth being arrested upon induced knockout of the *Nrde2-D174R* and *Nrde2Δ200* genes (Figure S1B). Thus, NRDE2 sustains cell growth independently of its protein interactors.

**Figure 1.**
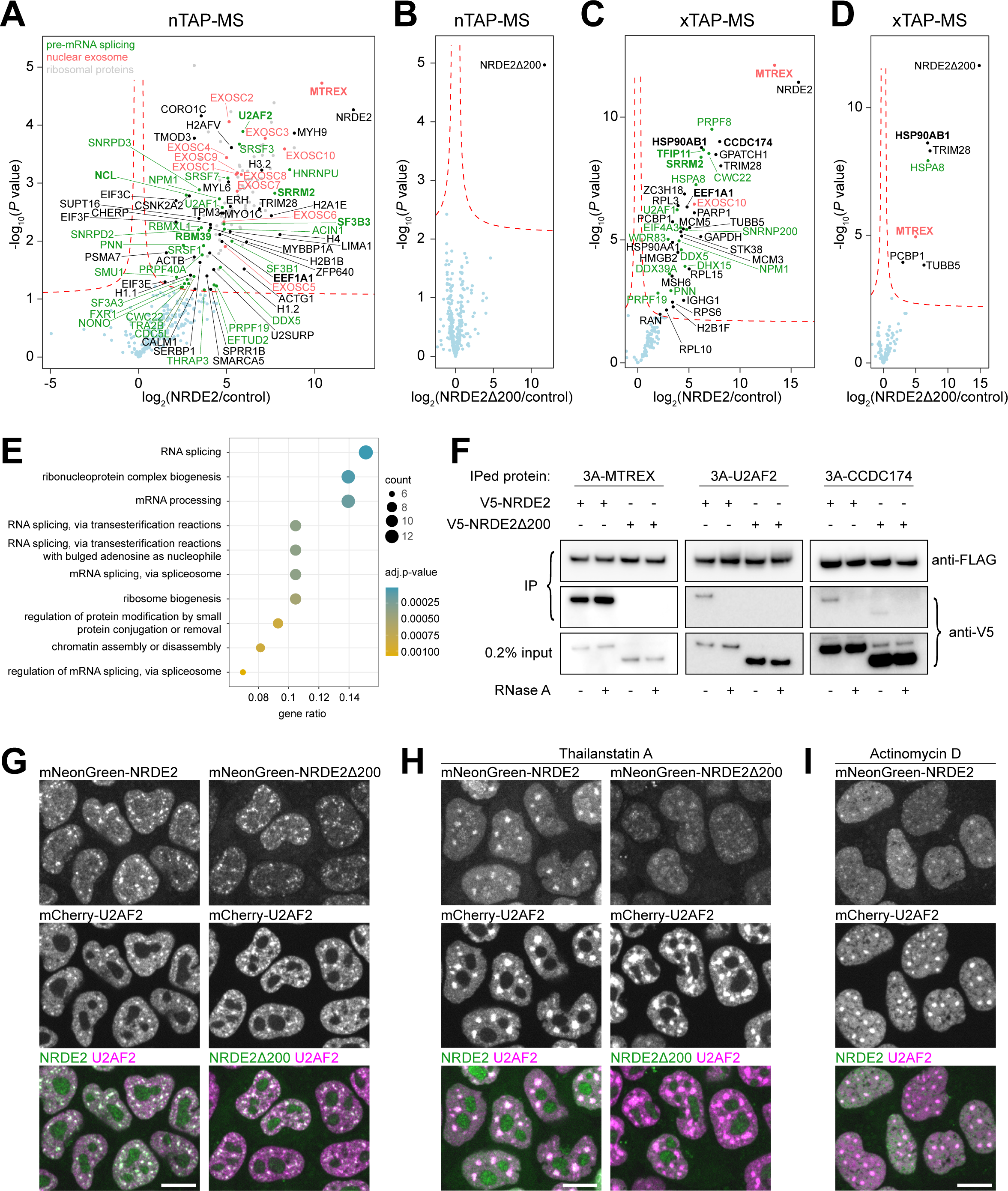
Protein interaction analysis links NRDE2 to pre-mRNA splicing. (A and B) Native TAP-MS analysis of 3A-NRDE2 (A) and 3A-NRDE2Δ200 (B) under low salt conditions (100mM NaCl) compared to untagged control and performed in three independent replicates for each sample. The red dashed line marks false discovery rate of 0.05. The components of the nuclear exosome are labeled in red. Proteins with GO term ’RNA splicing’ are depicted in green. Proteins highlighted in bold have also been identified in the yeast two-hybrid screen. Unlabeled grey dots represent significantly enriched ribosomal proteins. (C and D) Formaldehyde cross-linking TAP-MS analysis of 3A-NRDE2 (C) and 3A-NRDE2Δ200 (D) compared to untagged control and performed in three independent replicates for each sample. Same labeling scheme as in Figures 1A and 1B applies. (E) GO term analysis of 87 high-confidence hits from a yeast two-hybrid screen of full-length NRDE2 (See also Table S2). (F) Streptavidin pulldown and co-immunoprecipitation of endogenous 3A-tagged proteins from cells transiently overexpressing V5-NRDE2 or V5-NRDE2Δ200 fused to 2A-Puro in the presence or absence of RNase A. (G-I) Live-cell spinning-disk microscopy of endogenous mNeonGreen-tagged NRDE2 and NRDE2Δ200 with endogenous mCherry-tagged NS marker U2AF2 (CHUSAINOW et al., 2005). The imaging was performed in untreated cells (G), cells treated for 6 hours with 1 μM splicing inhibitor Thailanstatin A (H) or cells treated for 2 hours with 5 μg/ml transcription inhibitor Actinomycin D (I). Scale bar 10 μm.

### NRDE2 Localization to Nuclear Speckles Depends on Active pre-mRNA Splicing

NRDE2 has a relatively large PPI network. To identify proteins in direct proximity to NRDE2 *in vivo* we developed a limited cross-linking TAP-MS (xTAP-MS) protocol, which employs a short fixation with a low concentration formaldehyde followed by stringent lysis and washing conditions. NRDE2 xTAP-MS binding proteins were enriched for MTREX and splicing factors (Figure 1C), including proteins involved in different splicing steps, such as the DDX5 helicase, which regulates U1 snRNP-5’SS interactions (Liu, 2002), the U2AF1 component of the pre-spliceosome 3’ splice site-defining U2AF complex (Chen et al., 2017), the core spliceosome scaffold PRPF8 with 5’exon-stabilizing proteins SRRM2/SRM300 and CWC22, which are present through all stages of active spliceosome rearrangements (Zhang et al., 2018, 2019), as well as late spliceosome disassembly factors DHX15/PRP43 and TFIP11 (Yoshimoto et al., 2009). In addition, CCDC174 was also detected by xTAP-MS, indicating that it comes in proximity to NRDE2 in mammalian cells. NRDE2-D174R cross-linked to a similar set of proteins, including MTREX (Figure S1I), suggesting that NRDE2-D174R and MTREX still co-localize *in vivo* despite their reduced binding affinity. In agreement with the native pulldown, only a few mostly high-abundant proteins were detected in NRDE2Δ200 xTAP-MS along with low levels of MTREX, but with no spliceosome components present (Figure 1D).

The prevalence of splicing factors in the NRDE2 interactome was further confirmed by a yeast two-hybrid (Y2H) screen (Figure 1E), identifying MTREX and CCDC174 as high confidence interactors together with factors from different stages of spliceosome assembly, including U2AF2 and TFIP11 (Table S2). Selected Y2H interactions were validated by reciprocal co-IP. Unlike MTREX, NRDE2 interactions with U2AF2 and CCDC174 required an intact RNA component (Figure 1F), and a weak RNase-sensitive interaction was also maintained between CCDC174 and NRDE2Δ200.

Consistent with a splicing-related activity, fluorescent light microscopy revealed that NRDE2 was strongly enriched in nuclear splicing speckles (NSs) (Figure 1G). NSs are interchromatin granules with high concentration of splicing factors that have been linked to post-transcriptional processing of retained introns harboring weak splice sites (Galganski et al., 2017; Girard et al., 2012; Gordon et al., 2021; Vargas et al., 2011). NRDE2Δ200 showed a very similar subcellular distribution to wild type NRDE2, despite its lack of interactions with splicing factors (Figure 1G). Chemical inhibition of splicing with Thailanstatin A (Liu et al., 2013) resulted in the NRDE2Δ200 signal becoming more dispersed and NRDE2-D174R accumulated in nucleoli, with wild-type NRDE2 remaining concentrated in enlarged NSs (Figure 1H and S1J). In contrast, a transcriptional shut-off by Actinomycin D led to a substantial reduction of NS signal even for wild-type NRDE2 (Figure 1I). Overall, these results suggest that NRDE2 is actively recruited to and associates with pre-mRNAs that undergo splicing, independently of its interaction with splicing factors.

### NRDE2 and CCDC174 Bind to 5’ Splice Sites

To explore the RNA binding potential of NRDE2, we developed a Benzonase-assisted RNA cross-linking immunoprecipitation protocol coupled to high-throughput sequencing (BCLIP-seq) (Figure 2A)(Lee and Ule, 2018). Our approach offers a streamlined and sensitive alternative to existing CLIP techniques (Lee and Ule, 2018), enabling us to perform RNA interaction profiling for low abundant proteins such as NRDE2. The detergent-resistant Benzonase nuclease allows whole-cell lysis after UV cross-linking under highly denaturing conditions with simultaneous fragmentation of nucleic acids. The released protein-RNA cross-links are tandem affinity purified under stringent conditions, polyadenylated on beads, and the RNA is isolated for adaptor addition by a template switching-compatible reverse transcriptase (Turchinovich et al., 2014). This ligation-free cDNA library construction yields libraries of 20-150bp long inserts (Figure S2A).

**Figure 2.**
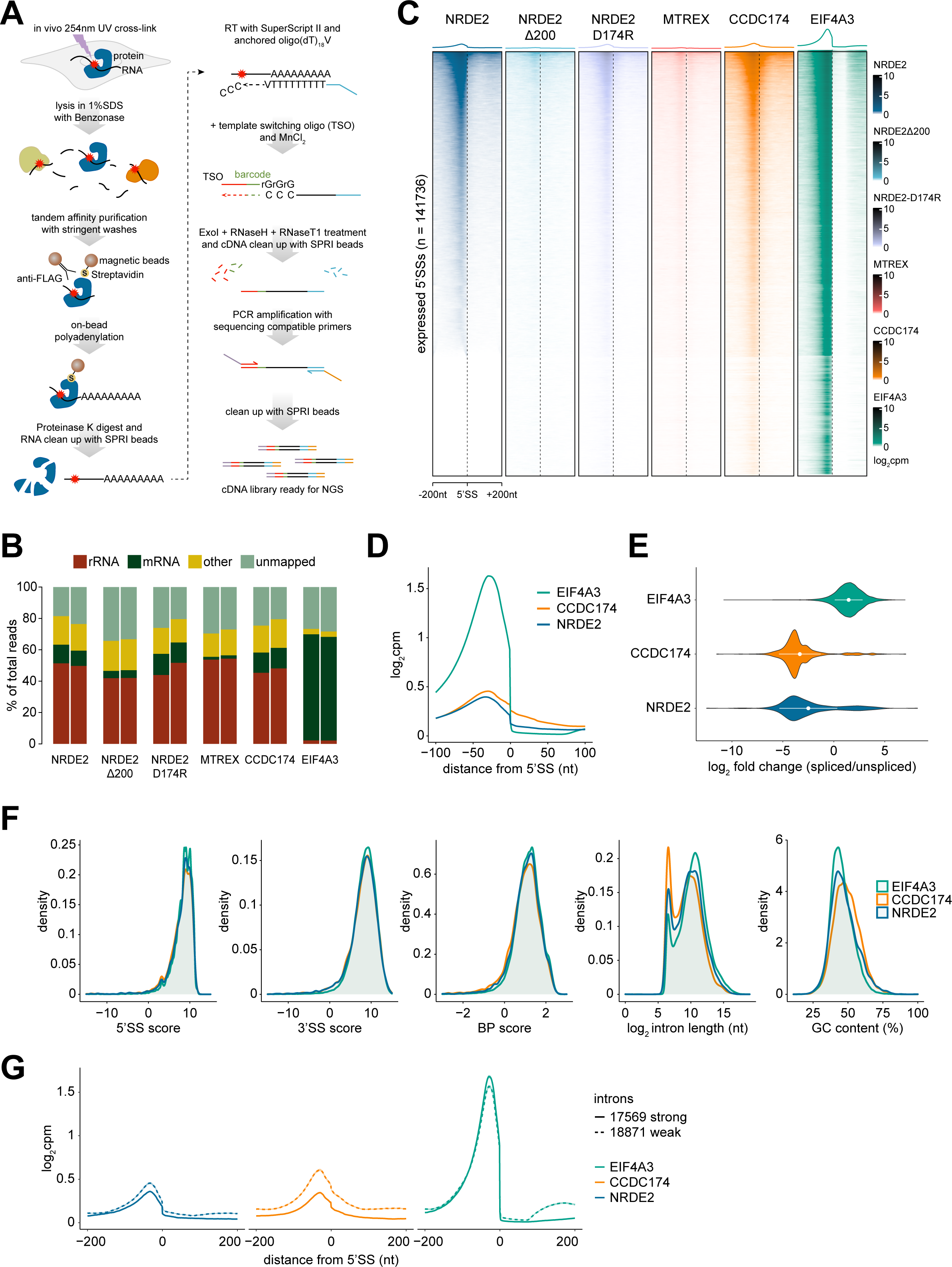
NRDE2 and CCDC174 bind to pre-mRNA 5’splice sites. (A) Schematic workflow of the BCLIP-seq protocol. TSO = Template Switching Oligo. (B) Mapping characteristics of the BCLIP-seq reads. The category ’rRNA’ contains reads mapping to mouse rDNA (GenBank: BK000964), category ’mRNA’ comprises reads mapping to protein coding mature mRNAs and category ’other’ includes all remaining reads mapping to the mouse genome. The two bars in each sample represent two independent BCLIP-seq replicates. (C) Heatmaps of the normalized BCLIP-seq signal intensity centered at annotated 5’SSs of all transcripts expressed in mESCs. Splice sites are ordered by decreasing NRDE2 signal. (D) Metaplots of normalized NRDE2, CCDC174 and EIF4A3 BCLIP-seq signal around all annotated expressed 5’SSs. (E) Violin plot showing ratios of spliced vs unspliced NRDE2, CCDC174 and EIF4A3 BCLIP-seq reads spanning annotated 5’SSs. Only 5’SSs of introns longer than 1 kb with a minimum of 10 splice site-overlapping reads were included in the analysis. White dots and bars represent the mean and standard deviation of all analyzed introns. (F) Density plots of the distribution of NRDE2, CCDC174 and EIF4A3 BCLIP-seq signal on all expressed introns ranked by their 5’SS strength, 3’SS strength, branch point strength, intron length and GC content. (G) Metaplots of normalized NRDE2, CCDC174 and EIF4A3 BCLIP-seq signal around 5’SSs of introns selected based on their 5’SS, 3’SS and branch point strength. Solid and dashed lines represent signal on introns with all three features above and below median strength, respectively.

We performed two BCLIP-seq replicates each for NRDE2, NRDE2Δ200, NRDE2-D174R, MTREX, and CCDC174. The exon junction complex (EJC) component EIF4A3 (Boehm and Gehring, 2016) acted as a positive control for a splicing-dependent RNA binding protein. The replicate samples correlated strongly (Figure S2B), which allowed us to merge the replicates into one dataset for most of the subsequent analyses. More than half of the mapping reads in all variant NRDE2, MTREX, and CCDC174 libraries matched rRNA (Figure 2B). MTREX was enriched at its known rRNA binding sites (Thoms et al., 2015), whereas no specific binding sites could be found for the NRDE2 variants, and CCDC174 was only mildly enriched in the 5’ external transcribed spacer (Figure S2C). The rRNA signal in NRDE2 and CCDC174 BCLIP-seq data could therefore represent non-specific binding to abundant RNA. However, the EIF4A3 data indicated that rRNA was not a common contaminant of the BCLIP method (Figure 2B). Therefore, we did not rule out potential functions of NRDE2 and CCDC174 in conjunction with rRNA .

Consistent with a potential role in pre-mRNA splicing, the majority of NRDE2 BCLIP peaks within protein-coding genes overlapped with exons and 5’SSs (Figure S2D). When considering all expressed 5’SSs, NRDE2 and, to a lesser extent, NRDE2-D174R were enriched at a subset of 5’SSs similar to CCDC174, whereas NRDE2Δ200 and MTREX showed very little binding around 5’SSs. EIF4A3 was enriched at most 5’SSs, as expected for an EJC factor (Figure 2C). NRDE2 and CCDC174 signals peaked 30nt upstream of 5’SSs, at the same position as EIF4A3, but, unlike EIF4A3, extended into introns. This was particularly prominent for CCDC174 (Figure 2D). Both NRDE2 and CCDC174 bound mostly to unspliced 5’SSs, implying that they associate with pre-mRNA before or during splicing (Figure 2E).

NRDE2- and CCDC174-bound 5’SSs showed a slight enrichment for shorter GC-rich introns, however, we could not distinguish them from control EIF4A3-bound 5’SSs by looking at the strength of individual intron-defining features, such as 5’SS, 3’SS, or branch point (Figure 2F), which were calculated using the Matt toolkit (Gohr and Irimia, 2018). Nevertheless, the overall NRDE2 and CCDC174 BCLIP signal was increased on 5’SSs of introns below median strength for all three features combined (Figure 2G). Thus, NRDE2 and CCDC174 are RNA-binding proteins that associate with unspliced 5’SSs.

### *Nrde2* and *Mtrex* Knockouts Induce a 2C-like State

RNA sequencing (RNA-seq) analysis of *Nrde2*-KO cells revealed a large group of differentially expressed genes (Figure 3A). NRDE2Δ200-expressing cells showed similar, but lower-magnitude expression changes, indicating that NRDE2 function is compromised but not abolished by the N-terminal truncation, consistent with the viability of *Nrde2Δ200* cells. Most transcripts downregulated in *Nrde2*-KO cells were also less abundant in CCDC174-depleted cells (*Ccdc174-dTAG*), whereas a large group of transcripts upregulated in *Nrde2*-KO cells were also upregulated upon conditional *Mtrex* knockout. These were highly enriched for genes defining a subpopulation of mESCs that resemble the gene-expression state of 2-cell (2C) embryos (Macfarlan et al., 2012) (Figure 3B). The expression of 2C genes correlates with the activity of MERVL, a family of retrotransposons whose expression in mESCs is regulated by facultative heterochromatin (fHC). The NRDE2 homologs in *C. elegans* and *S. pombe* have been linked with fHC regulation (Guang et al., 2010; Lee et al., 2013) and MERVL and its long terminal repeat (LTR) promoter, MT2_Mm, were the most upregulated repetitive elements in both *Nrde2-*KO and *Mtrex-*KO cells (Figure S3A).

**Figure 3.**
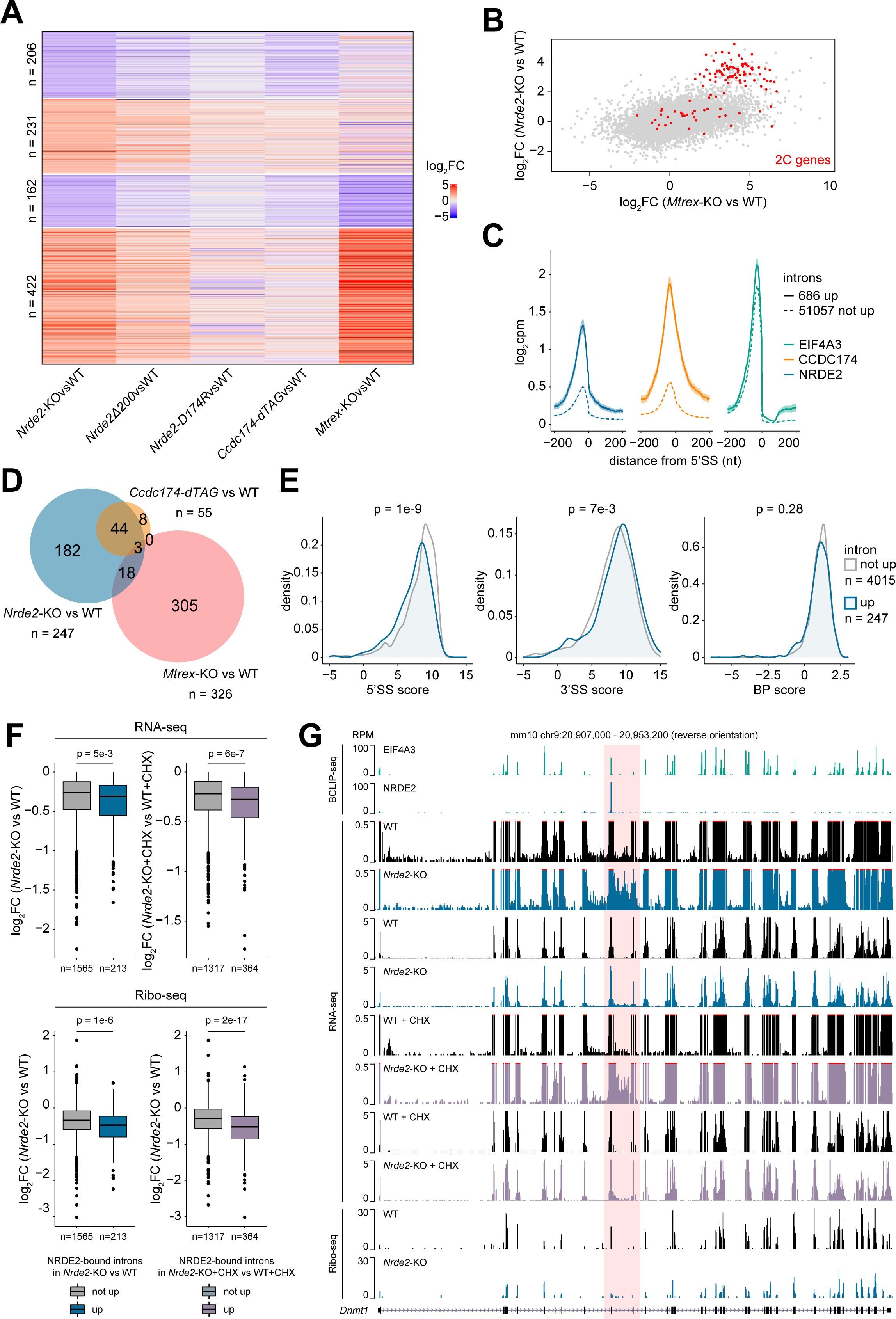
NRDE2 regulates splicing of introns with weak 5’splice sites. (A) Differential gene expression based on duplicate RNA-seq analysis of *Nrde2*-KO (4 days of 0.1 μM 4OHT treatment), *Nrde2Δ200*, *Nrde2-D174R*, *Ccdc174-dTAG* (24 hours of 0.5 μM dTAG-13 treatment), and *Mtrex*-KO (3 days of 0.1 μM 4OHT treatment) cells compared to the matching untreated and wild-type controls (WT). Heatmap consists of genes differentially expressed (fold change > 2, adjusted p-value < 0.01) in *Nrde2*-KO cells. Genes are divided into four clusters depending on expression changes in *Nrde2*-KO and *Mtrex*-KO samples. (B) Scatter plot comparing differential gene expression in *Nrde2*-KO and *Mtrex*-KO cells relative to the corresponding WT cells. Each dot represents a single gene. Genes upregulated in 2C-like cells (Macfarlan et al., 2012) are highlighted in red. (C) Metaplots of normalized NRDE2, CCDC174 and EIF4A3 BCLIP-seq signal around 5’SSs of introns upregulated (solid line) or not upregulated (dashed line) in *Nrde2*-KO cells compared to WT. (D) Overlap of NRDE2 target introns (introns with a NRDE2 BCLIP-seq peak at their 5’SS) upregulated in *Nrde2*-KO, *Ccdc174-dTAG* or *Mtrex*-KO cells. (E) Density plots of the distribution of NRDE2 target introns upregulated (blue line) or not upregulated (grey line) in *Nrde2*-KO cells based on their 5’SS, 3’SS and branch point strength. P-values were calculated using Wilcoxon rank-sum test. (F) Differential expression (RNA-seq) of genes containing NRDE2 target introns in *Nrde2*-KO cells (4 days of 0.1 μM 4OHT treatment) untreated (left) or treated (right) for 4 hours with 100 μg/ml cycloheximide (CHX). Genes are split into two groups: those containing only target introns not upregulated (grey) and those containing at least one target intron upregulated in *Nrde2*-KO cells untreated (blue) or treated with CHX (purple). Bottom boxplots show differential ribosome occupancy for the same genes in *Nrde2*-KO cells relative to WT. P-values were calculated using Wilcoxon rank-sum test. (G) UCSC genome browser screenshot showing normalized BCLIP-seq, RNA-seq and Ribo-seq tracks over *Dnmt1* gene locus. Position of the NRDE2 target intron 10 is highlighted in red.

To test the hypothesis that NRDE2 acts with MTREX to negatively regulate MERVL expression, we created cells with an MT2_Mm-driven mNeonGreen (2C reporter) inserted in a genomic locus replacing an active MERVL (Figure S3B). When combined with knockouts of fHC-regulating enzymes G9A and KDM1A (Macfarlan et al., 2011, 2012), *Nrde2* knockout had an additive effect on MERVL and 2C gene upregulation, as a consequence of the increased ratio of 2C reporter-positive cells (Figure S3C and S3D). These results indicate that NRDE2 influences MERVL expression independently of fHC formation. NRDE2 could be regulating MERVL at a post-transcriptional level, given its link to MTREX and the nuclear exosome. However, reducing the MT2_Mm reporter to shorter variants that preserve promoter activity but ultimately lead to reporter transcripts devoid of MERVL sequences did not alleviate NRDE2 dependence (Figure S3E and S3F). Therefore, the observed upregulation of MERVL and 2C genes upon *Nrde2* or *Mtrex* knockout likely represents a pleiotropic effect caused by altered RNA metabolism, rather than an escape from the specific action of NRDE2 and MTREX on these genes. This conclusion is further supported by negligible effects of the NRDE2-D174R mutation on gene expression, including 2C genes (Figures 3A and S3G).

### NRDE2 and CCDC174 Regulate Weak 5’ Splice Sites

Given NRDE2 and its interactor CCDC174 bind to 5’SSs, we hypothesized that pre-mRNA splicing could be defective in mutant cell lines, manifest as globally increased intronic signals in the RNA-seq data. To exclude transcription activation effects, we analyzed downregulated transcripts only (log_2_ fold change < 0). In both *Nrde2-*KO and *Ccdc174-dTAG* cells, we found hundreds of misspliced upregulated introns that were generally enriched for NRDE2 and CCDC174 binding at their 5’SSs (Figures 3C and S4A). However, the upregulated introns constituted only a small fraction of all the introns defined as NRDE2 and CCDC174 target introns (5.8% and 2%, respectively), for which the peak-calling analysis of the BCLIP-seq data identified a NRDE2 or CCDC174 peak at their 5’SS. The majority of the target introns with NRDE2- or CCDC174-bound 5’SSs remained unaffected. Misspliced target introns largely overlapped in the *Nrde2-*KO and *Ccdc174-dTAG* data sets, but they were distinct from target introns upregulated in *Mtrex-*KO cells (Figures 3D and S4B). Hence, ablation of NRDE2 or CCDC174 can have a direct effect on splicing of a subset of introns they associate with, which is not observed for MTREX. Therefore, NRDE2 and CCDC174 likely function independently of MTREX in pre-mRNA splicing. We note that the number of NRDE2-bound upregulated introns was much smaller in *Nrde2Δ200* compared to *Nrde2*-KO cells, suggesting the N-terminally truncated NRDE2 functions as a hypomorph (Figure S4C).

Our observation that splicing of only a minor fraction of NRDE2 and CCDC174-bound introns was affected in *Nrde2-*KO cells could be due in part to the limited sensitivity of RNA-seq for detecting unstable misspliced transcripts. Blocking translation-dependent RNA surveillance by cycloheximide (CHX) treatment led to an increase in misspliced introns (12% of NRDE2- bound and 17% of CCDC174-bound) that overlapped between different conditions (Figure S4C). However, the majority of introns remained unchanged even after CHX treatment, prompting us to investigate the selective sensitivity of target introns to NRDE2 or CCDC174 depletion. Analysis of intron features revealed that NRDE2-bound introns upregulated in *Nrde2-*KO cells possess weaker 5’SSs, while their 3’SSs and branch points are comparable to non-affected introns (Figure 3E). The difference in 5’SS strength was more pronounced in the group of introns upregulated in CHX-treated *Nrde2*-KO cells. The same was observed for *Ccdc174*-bound introns (Figure S4D and S4E).

The aberrant splicing of introns in *Nrde2-*KO and *Ccdc174-dTAG* cells resulted in an overall decrease of the respective mRNA signal (Figures 3F and S4F). However, the impact on mRNA levels was generally small, possibly due to the stability of the aberrantly spliced transcripts. To assess the effect on translation, we performed ribosome profiling (Ribo-seq) in wild-type and *Nrde2-*KO cells. Compared to RNA-seq, a larger Ribo-seq signal reduction was observed for messages whose splicing was affected by *Nrde2* knockout (Figure 3F). For example, intron 10 of the *Dnmt1* pre-mRNA has a suboptimal 5’SS that is bound by NRDE2 and CCDC174. Intron 10 levels were upregulated in *Nrde2-*KO and CCDC174-depleted cells without a substantial effect on mRNA levels, whereas ribosome occupancy was markedly reduced (Figures 3G and S4G). Thus, overall, NRDE2 and CCDC174 are required for efficient splicing of a largely overlapping set of target introns with suboptimal 5’SS sequences.

### NRDE2 Is Necessary for Correct Splice Site Choice

Defective splicing can lead to intron retention, splicing from a cryptic 5’SS, or splicing to a cryptic 3’SS. Analysis of novel splicing events in upregulated NRDE2-bound introns revealed frequent cryptic 5’SS usage upon *Nrde2* knockout (Figure 4A). Likewise, CCDC174-bound upregulated introns showed higher incidence of cryptic 5’SS usage upon CCDC174 depletion (Figure S5A). The first intron of the essential cell cycle regulator gene *Cdk2*, which contains a weak 5’SS strongly bound by NRDE2 and CCDC174, provides a representative example of such aberrant splice site choice. Examination of our RNA-seq data uncovered a shift in *Cdk2* intron 1 splicing from the annotated 5’SS to several cryptic 5’SSs in cells lacking NRDE2 or CCDC174 (Figure 4B). Splicing from the cryptic 5’SSs generated non-functional transcripts because of an in-frame stop codon immediately downstream of the first annotated 5’SS, resulting in a substantial reduction in Ribo-seq signal (Figure 4B). Usage of multiple alternative cryptic 5’SSs was confirmed by RT-PCR with primers binding to *Cdk2* exons 1 and 3 (Figure 4C). We observed a similar but weaker aberrant splicing pattern in NRDE2Δ200-expressing cells. Usage of a small number of cryptic 5’SSs was also detected in *Mtrex-*KO cells, likely due to compromised NRDE2 stability. In contrast, depletion of the general splicing factors SmE (a component of the snRNP-stabilizing Sm ring complex) or CWC22 resulted in minimal cryptic 5’SS usage. Notably, all the alternative 5’SSs in the *Cdk2* intron 1 that are utilized in the absence of NRDE2 or CCDC174 consist of suboptimal sequences. Thus, NRDE2 and CCDC174 appear to be dispensable for general splicing, but are required for appropriate splice site choice.

**Figure 4.**
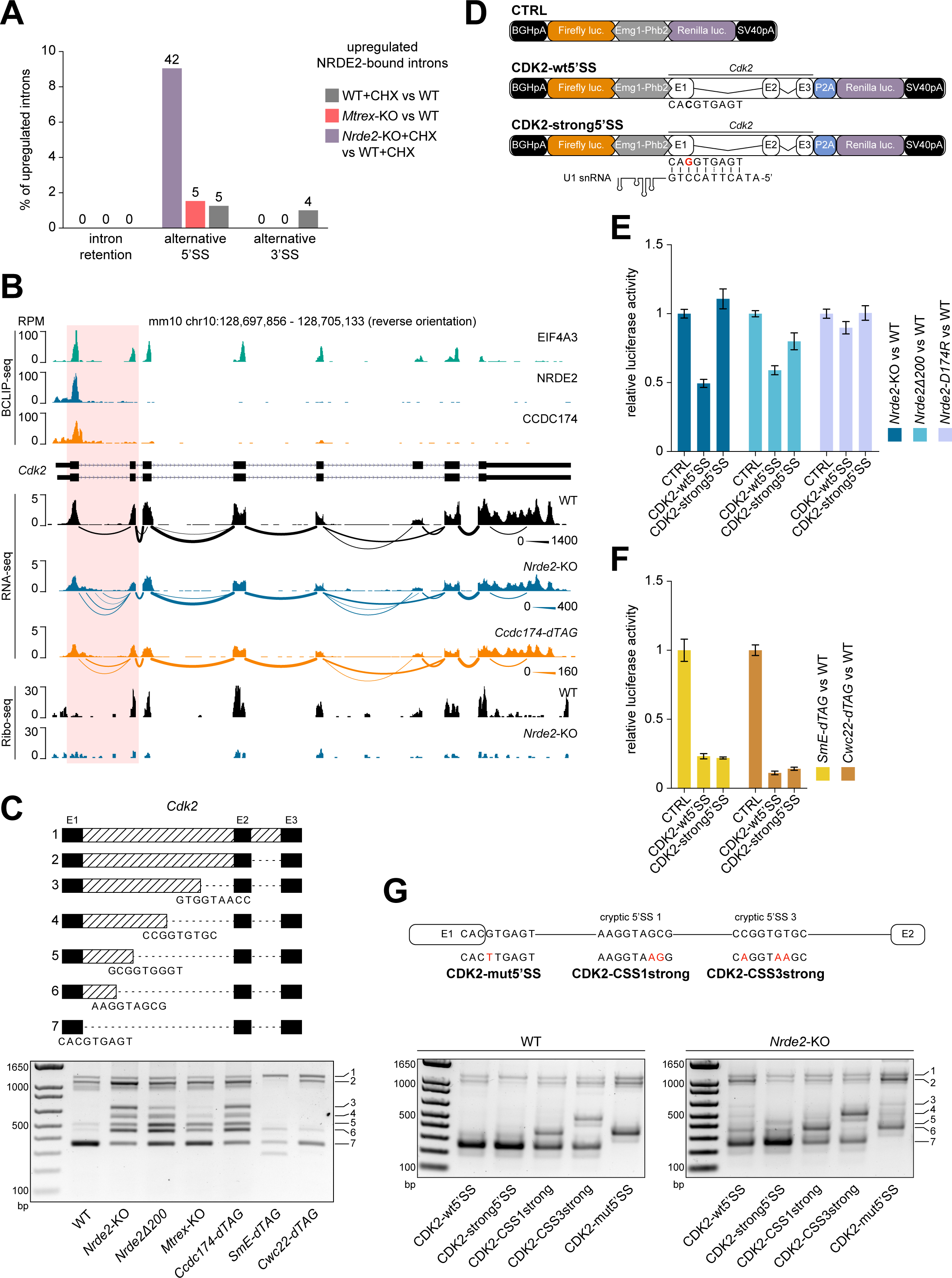
NRDE2 regulates 5’SS selection and splicing of its target gene Cdk2. (A) rMATS analysis (Shen et al., 2014) of alternative splicing events within NRDE2 target introns upregulated in *Mtrex*-KO cells and in CHX-treated WT and *Nrde2*-KO cells relative to their corresponding WT and untreated controls. (B) UCSC genome browser screenshot showing normalized BCLIP-seq, RNA-seq and Ribo-seq tracks over *Cdk2* gene locus combined with a Sashimi plot of *Cdk2* splicing patterns detected in the RNA-seq data. The thickness of the splice junction connecting lines reflects the number of corresponding spliced RNA-seq reads with a scale provided on the right. NRDE2 and CCDC174 target intron 1 is highlighted in red. (C) RT-PCR analysis of *Cdk2* intron 1 splicing with primers binding in exon 1 and 3. The additional bands in *Nrde2*-KO sample were sequenced and the identified cryptic 5’SSs are depicted in the scheme on top. (D) Schematic overview of the CDK2 dual luciferase reporters. The point mutation introduced in *Cdk2* intron 1 5’SS to improve U1 snRNA complementarity in the CDK2-strong5’SS reporter is depicted in red. (E and F) Dual luciferase assay of the transiently transfected CDK2 reporters. Plotted is the average value of normalized Renilla luciferase activity from three independent experiments each performed in triplicate transfections. Error bars represent standard deviation. (G) RT-PCR analysis of CDK2 reporter splicing in transiently transfected WT and *Nrde2*-KO cells with primers binding in *Cdk2* exon1 and in the P2A peptide region. The additional mutations introduced in the CDK2-wt5’SS reporter are depicted in red. The band numbering on the right corresponds to the scheme in panel (C).

To determine whether NRDE2 splice site choice is modulated by 5’SS strength and to validate *Cdk2* intron 1 as a NRDE2 target, we fused *Cdk2* exons 1 - 3 with the *Renilla* luciferase gene in a dual luciferase reporter plasmid (CDK2-wt5’SS). We also generated a CDK2 reporter construct in which the sequence of the first 5’SS was optimized by a single point mutation (CDK2-strong5’SS), improving U1 snRNA complementarity (Figure 4D). Compared to wild-type cells, the activity of CDK2-wt5’SS was 2-fold lower when expressed in *Nrde2-*KO cells whereas CDK2-strong5’SS was insensitive to NRDE2 deficiency (Figure 4E). Thus, 5’SS strength is a predictive feature of NRDE2-dependent pre-mRNA splicing.

CDK2-wt5’SS activity was insensitive to the NRDE2-D174R mutation, but was affected by the N-terminal truncation (NRDE2Δ200) (Figure 4E), further supporting the idea that NRDE2 functions independently of MTREX in pre-mRNA splicing, and that the N-terminus of NRDE2 contributes to its splicing related activity. In comparison to *Nrde2-*KO cells, reduction of CDK2 reporter activity was more substantial in cells lacking SmE or CWC22, regardless of 5’SS strength (Figure 4F), indicating that NRDE2 is not an obligate splicing factor, but is necessary for the efficient use of weak 5’SSs. To support and generalize this conclusion, we generated reporter constructs containing NRDE2-bound sequences of the DNA damage response regulator *Tti1* gene, where NRDE2 and CCDC174 bind the 5’SS of intron 5 (Figure S5B and S5C). A two-nucleotide substitution that strengthened the weak 5’SS of intron 5 abolished the NRDE2-dependency of *Tti1* splicing (Figures S5D - S5F), consistent with the CDK2 reporters. The first *Cdk2* intron harbors multiple cryptic 5’SSs, which allowed us to dissect the 5’SS choice hierarchy. We introduced several mutations into the CDK2-wt5’SS luciferase reporter: mutations that improve the strength of either the first or third cryptic 5’SS, or a mutation that inactivates the annotated 5’SS (Figure 4G). As predicted by our model, lack of NRDE2 resulted in usage of four additional cryptic 5’SSs located downstream of the annotated 5’SS. The shift in splice site choice upon *Nrde2* knockout was diminished if the annotated 5’SS was mutated to a strong 5’SS. Increasing the strength of the first or third cryptic 5’SS resulted in their usage in wild-type cells, along with the annotated 5’SS. This balance shifted towards the strengthened cryptic 5’SSs in *Nrde2-*KO cells, indicating that NRDE2 enhances splicing from the most upstream weak 5’SS, even when followed by a stronger downstream 5’SS. Notably, inactivation of the annotated 5’SS redirected NRDE2 regulation to the neighboring downstream cryptic 5’SS, which was the exclusive 5’SS used in wild-type cells. In the absence of NRDE2, the remaining cryptic sites were chosen again (Figure 4G). Thus, NRDE2 promotes splicing from the most upstream of a series of 5’SSs.

### CCDC174 Function Depends on NRDE2

NRDE2 and CCDC174 promote splicing at many of the same weak 5’SSs, with, for example, the activity of the CDK2 and TTI1 luciferase reporters being the same in *Nrde2-*KO and *Ccdc174-dTAG* cells (Figure 5A). Simultaneous depletion of NRDE2 and CCDC174 had no additive effect on reporter activity, suggesting that the two proteins act together. This was further supported by xTAP-MS of CCDC174, which identified a set of splicing factors similar to those present in the NRDE2 interactome (Figures 5B and 1C). CCDC174 also accumulated in NSs, and its accumulation was reduced by *Nrde2* knockout, whereas NRDE2 NS localization remained unchanged in CCDC174-depleted cells (Figure 5C). Hence, NRDE2 recruits CCDC174 to NSs to exert its splicing-related function.

**Figure 5.**
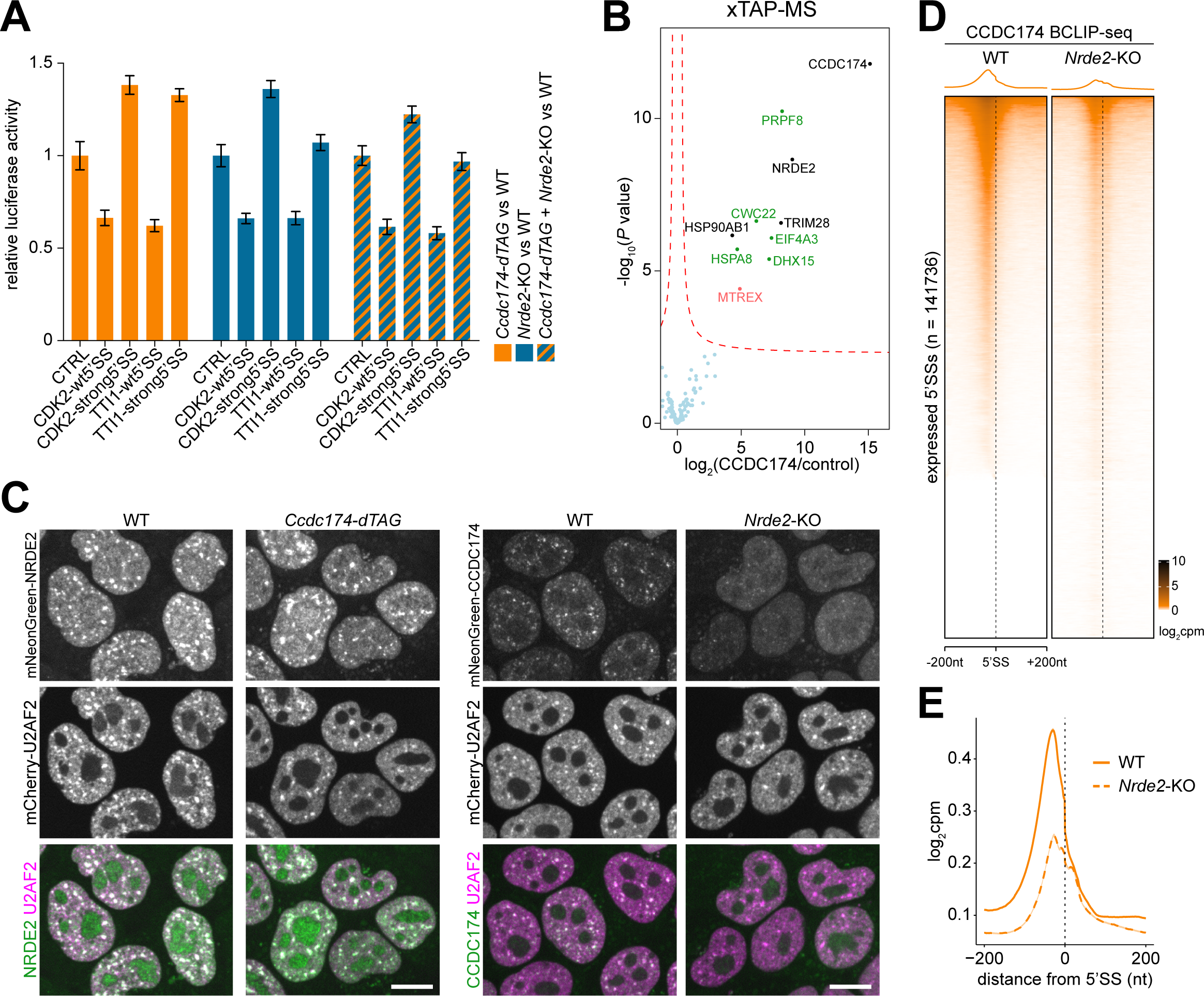
NRDE2 is required for proper recruitment of CCDC174 to 5’SSs. (A) Dual luciferase assay of the transiently transfected CDK2 and TTI1 reporters in cells allowing simultaneous depletion of NRDE2 and CCDC174. Plotted is the average value of normalized Renilla luciferase activity from three independent experiments each performed in triplicate transfections. Error bars represent standard deviation. (B) Formaldehyde cross-linking TAP-MS analysis of 3A-CCDC174 compared to untagged control and performed in three independent replicates for each sample. Same labeling scheme as in Figures 1A and 1B applies. (C) Live-cell spinning-disk microscopy of endogenous mNeonGreen-tagged NRDE2 and CCDC174 with endogenous mCherry-tagged U2AF2 in cells allowing depletion of CCDC174 and NRDE2, respectively. Scale bar 10 μm. (D) Heatmaps of the normalized CCDC174 BCLIP-seq signal intensity in WT and *Nrde2*-KO cells centered at annotated 5’SSs of all transcripts expressed in mESCs. Splice sites are ordered by decreasing CCDC174 signal in the WT sample. (E) Metaplots of normalized CCDC174 BCLIP-seq signal around all annotated expressed 5’SSs in WT (solid line - same data as in Figure 2D) and *Nrde2*-KO (dashed line).

To analyze the relationship between NRDE2 and CCDC174, we performed CCDC174 BCLIP-seq in *Nrde2-*KO cells. Overall binding of CCDC174 at 5’SSs was substantially reduced in the absence of NRDE2 (Figure 5D). Interestingly, lack of NRDE2 mainly affected CCDC174 binding in the exonic region upstream of 5’SSs, whereas the average low-level intronic CCDC174 BCLIP-seq signal remained comparable between wild-type and *Nrde2*-KO cells (Figure 5E). Residual CCDC174 binding to 5’SSs could therefore persist in the absence of NRDE2, yet at a different preferred position. Taken together, these results demonstrate that NRDE2 is required for efficient recruitment of CCDC174 to NSs and association with 5’SSs, where the two proteins cooperate to sustain weak 5’SS selection and pre-mRNA splicing.

### NRDE2 and CCDC174 Bind to U1 snRNA

Participation of NRDE2 in 5’SS choice raises the question how 5’SSs are recognized by NRDE2. Furthermore, how does NRDE2 recruit CCDC174, given that they interact weakly in an RNA-dependent manner? Considering NRDE2Δ200 localized to NSs and was partially functional, despite losing interactions with splicing factors and no longer cross-linking to pre-mRNAs, it seemed unlikely that NRDE2 engages with the spliceosome via interaction with one of its proteinaceous components or by binding to a specific sequence motif in pre-mRNA. Instead, the NRDE2 BCLIP-seq data sets were highly enriched for reads mapping to major spliceosome snRNAs (ms-snRNAs), an association that is highly specific, given we did not observe this for MTREX or EIF4A3 (Figure 6A). The portion of ms-snRNA mapping reads further increased in NRDE2-D174R libraries and reached maximal levels in NRDE2Δ200 samples (5% of total reads). CCDC174 was equally strongly associated with ms-snRNAs (4% of total reads). No appreciable enrichment in signal was observed for minor spliceosome snRNAs. The majority of ms-snRNA reads in all NRDE2 and CCDC174 samples mapped to U1 snRNA, from nearly 60% for NRDE2 to more than 80% for NRDE2Δ200 and CCDC174 (Figure 6B). The association of NRDE2 and CCDC174 with U1 snRNA was confirmed under native conditions by immunoprecipitation followed by Northern blotting (Figure 6C). In addition, both NRDE2 and CCDC174 co-immunoprecipitated with an anti-m3G antibody, which recognizes the trimethylated guanosine cap of snRNAs, in an RNase-sensitive manner (Figure S6A). The BCLIP-seq signal covered most of the highly structured U1 snRNA sequence, with the highest enrichment over the stem-loops II and III (SL2 and SL3) for both NRDE2 and CCDC174 (Figure 6D), indicating that NRDE2 and CCDC174 contact U1 snRNA at multiple positions. In comparison, the two other most represented ms-snRNAs, U4 and U6, appeared to cross-link at distinct locations overlapping the U4/U6 stem III region (Figure S6B). U1 stem-loops II and III and U4/U6 stem III are in close proximity in the pre-catalytic spliceosomal pre-B complex (Figure S6C) (Charenton et al., 2019), suggesting how U1 snRNA-bound NRDE2 and CCDC174 might contact U4 and U6 snRNAs. Together, these results reveal that U1 snRNA is the primary RNA substrate of NRDE2. Given CCDC174 association with U1 snRNA was reduced in the absence of NRDE2 (Figure 6E), we conclude that CCDC174 preferably binds NRDE2-bound U1 snRNAs.

**Figure 6.**
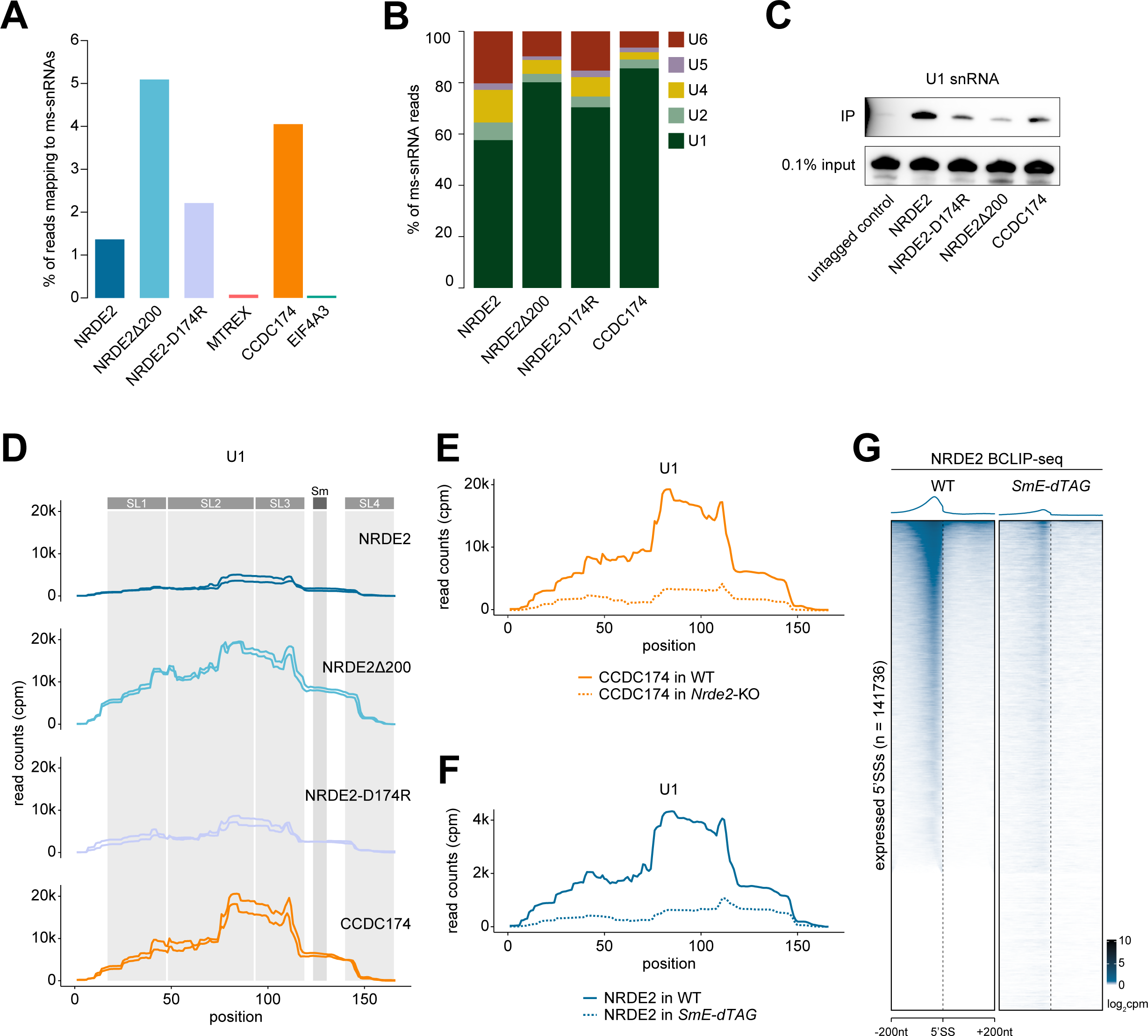
NRDE2 and CCDC174 bind to U1 snRNA. (A) Proportion of BCLIP-seq reads mapping to ms-snRNAs. (B) Relative distribution of ms-snRNA-mapping BCLIP-seq reads among individual snRNAs. (C) Northern blotting analysis of U1 snRNA in immunoprecipitates of tandem affinity purified control untagged cells and indicated endogenously 3A-tagged cells. (D) BCLIP-seq signal coverage over U1 snRNA sequence (including reads mapping to U1a1, U1b1 and U1b6 isoforms) showing the two BCLIP-seq replicates of each sample separately. Grey columns indicate the positions of the Sm ring binding site and the four stem-loops within the U1 snRNA secondary structure. (E) CCDC174 BCLIP-seq signal coverage over U1 snRNA sequence (including reads mapping to U1a1, U1b1 and U1b6 isoforms) in WT and *Nrde2*-KO cells. (F) NRDE2 BCLIP-seq signal coverage over U1 snRNA sequence (including reads mapping to U1a1, U1b1 and U1b6 isoforms) in WT and *SmE-dTAG* cells. (G) Heatmaps of the normalized NRDE2 BCLIP-seq signal intensity in WT and *SmE-dTAG* cells centered at annotated 5’SSs of all transcripts expressed in mESCs. Splice sites are ordered by decreasing NRDE2 signal in the WT sample which is the same as in Figure 2C.

The indiscriminate association of NRDE2 and CCDC174 with 5’SSs is explained by U1 snRNA-mediated targeting. To further test this model, we decided to perturb the structure of the Sm ring complex, which is necessary for U1 snRNA maturation and trafficking (Matera and Wang, 2014). Depletion of the Sm ring component SmE led to a significant reduction of NRDE2 BCLIP-seq signal on U1 snRNA (Figure 6F), demonstrating that NRDE2 association with U1 snRNA depends on a functional U1 snRNP assembly pathway. Consistent with U1 snRNA-mediated targeting of NRDE2 to pre-mRNAs, NRDE2 association with 5’SSs was substantially reduced upon SmE depletion (Figure 6G). Therefore, we propose a model in which NRDE2 and CCDC174 are directly recruited by U1-snRNA to sustain splicing of weak 5’SSs.

## DISCUSSION

Mammalian cells must cope with the challenge of selecting and splicing 5’SSs with suboptimal U1 snRNA complementarity. Mechanistic studies on weak 5’SS splicing have thus far focused on auxiliary splicing factors, mainly from the SR-protein family, that bind exonic splicing enhancer (ESE) motifs in the vicinity of 5’SSs to support the binding of canonical U1 snRNPs (Kohtz et al., 1994; Long and Caceres, 2008; Wu and Maniatis, 1993). Our study uncovers an alternative strategy that involves the conserved proteins NRDE2 and CCDC174 interacting directly with U1 snRNA to promote selection and splicing of weak 5’SSs.

### NRDE2 and CCDC174 Bind RNA

The composition of the NRDE2 and CCDC174 PPI networks links these two proteins to RNA metabolism, raising the question whether they have intrinsic RNA-binding activities. RNA binding is commonly probed using CLIP, which enriches for RNAs in direct contact with the protein of interest due to zero distance UV cross-linking (Lee and Ule, 2018). When using existing CLIP protocols for NRDE2 and unrelated low abundant proteins, we suffered from undesirable sample loss. Speculating that we might lose our samples because of the low UV cross-linking efficiency (Hafner et al., 2021) and during the many steps required to purify the RNA-protein complexes, we designed a simplified BCLIP-seq protocol that utilizes high affinity 3A tag components. This substantially reduced length and complexity of the cross-linked protein-RNA purification. The protocol is further streamlined by a ligation-free sequencing library construction using the template-switching activity of SuperScript II reverse transcriptase. Because this relies on the presence of Mn^2+^ ions, the resulting library might lack the single nucleotide resolution of the UV cross-linked nucleotide as suggested previously (Nostrand et al., 2017). Although neither NRDE2 nor CCDC174 contain predicted RNA-binding domains, our BCLIP-seq results revealed that both of them bind to RNA. Thus, NRDE2 and CCDC174 belong to a group of proteins whose amino-acid sequences are not predictive of RNA-binding activity (Albihlal and Gerber, 2018).

### NRDE2 and CCDC174 modulate weak 5’SSs

Our results revealed that the N-terminus of NRDE2 is dispensable for U1 snRNA binding, but is necessary for associating with pre-mRNAs at 5’SSs. This highlights U1 snRNA as the primary RNA substrate of NRDE2 and offers insights into the potential mechanisms that stabilize the association of the U1 snRNP with weak 5’SSs. It is possible that the N-terminus of NRDE2 mediates stabilizing interactions within the spliceosome once bound to the 5’SS. The N-terminus could also interact with the pre-mRNA, further stabilizing U1 snRNA association with weak 5’SS. Additional work will be required to fully elucidate these mechanisms, but such model may help explain the hypomorphic phenotype of the NRDE2Δ200 truncation.

Further, the multiple NRDE2 and NRDE2Δ200 contact sites on U1 snRNA suggest that a large portion of NRDE2 is dedicated to U1 snRNA binding. This leads to the provocative idea that NRDE2 could potentially mediate the formation of a non-canonical U1 snRNP. That NRDE2 is not co-purifying with canonical U1 snRNP-specific components U1A, U1C, and U1-70K supports such hypothesis (Kondo et al., 2015) (Figures 1A-1D). Our results further suggest that such an snRNP would create a scaffold for CCDC174 recruitment, generating a fully functional non-canonical U1 snRNP for stable binding and effective splicing of weak 5’SSs.

The above described model would explain why NRDE2 and NRDE2Δ200 interact with CCDC174 in an RNA-dependent manner, and is consistent with the shifted CCDC174 BCLIP-seq profile in *Nrde2*-KO cells, as residual CCDC174 association with canonical U1 snRNPs might lead to CCDC174 positioning close to the intronic part of 5’SSs, as opposed to CCDC174 binding to the upstream exonic region in the presence of NRDE2. Furthermore, it is possible that NRDE2 and CCDC174 potentially contact all 5’SSs but are mostly detected on weakly spliced introns because these likely have slower splicing kinetics, resulting in longer residence time and a higher chance of cross-linking. Consequently, only a minority of bound 5’SSs, those with inferior U1 snRNA complementarity, respond to loss of NRDE2 or CCDC174, whereas the rest are efficiently spliced by the canonical U1 snRNP.

Our results are also relevant for understanding splice-site choice. The luciferase reporter experiments revealed that NRDE2 stimulates splicing from the most upstream 5’SS of an intron, reminiscent of a previously described function of hnRNPA1 in *in vitro* splicing reactions (Mayeda and Krainer, 1992). In a co-transcriptional splicing model, this could simply be explained by the fact that the most upstream 5’SS emerges from the RNA polymerase first. Alternatively, it could be mediated by proteins binding to upstream ESEs. Indeed, we found several ESE-binding SR proteins interacting with NRDE2, but not with NRDE2Δ200 (Figures 1A and 1B) for which the most upstream 5’SS preference is partially lost. The luciferase reporter assays also revealed that NRDE2 participates in the splicing of target introns, rather than recruiting MTREX to degrade unspliced or incorrectly spliced transcripts, as suggested for the NRDE2 homolog in *S. pombe* (Zhou et al., 2015). Therefore, the function of the NRDE2-MTREX interaction in mammalian cells remains elusive. MTREX is required for NRDE2 stability, and based on our xTAP-MS results, it retains contact with NRDE2-D174R and NRDE2Δ200. Given its RNA helicase activity, a possible MTREX function could be unwinding of the NRDE2-bound, highly structured U1 snRNA. In this model, the arch interacting motif within the NRDE2 N-terminus would stabilize MTREX association to allow complete RNA unwinding.

### NRDE2 and CCDC174 Are Part of the Major Spliceosome

Our proteomic data and the BCLIP-seq signals at unspliced 5’SSs indicate that NRDE2 and CCDC174 associate with spliceosome components during splicing. However, they have not been identified in spliceosome composition studies (Rappsilber et al., 2002; Wahl et al., 2009; Zhou et al., 2002). Such analyses, both at the proteomic and structural level, have relied on spliceosomes pre-assembled on model pre-mRNA substrates with optimal splicing features, suggesting that they may not reflect the full scope of *in vivo* spliceosome diversity. In structural studies, the spliceosome behaves as a dynamic assembly progressing through distinct pre-catalytic, catalytic, post-catalytic and disassembly steps accompanied by considerable rearrangements, including U1 snRNP dissociation before the catalytic step (Wilkinson et al., 2019). Our data demonstrate simultaneous NRDE2 and CCDC174 association with the pre-catalytic spliceosome, represented by binding to U1 snRNA and unspliced 5’SSs, and with the late post-catalytic spliceosome via interactions with disassembly factors DHX15 and TFIP11. Hence, we speculate that NRDE2 and CCDC174 may be involved in an alternative spliceosome assembly pathway.

Notably, NRDE2 interactions with splicing factors were revealed by nTAP-MS only after digesting the samples with Benzonase nuclease, which loosens large RNP assemblies. Moreover, core spliceosome components and disassembly factors, as well as CCDC174, only appeared in xTAP-MS under harsh denaturing conditions, and freeze-thawing the purified xTAP-MS samples on magnetic beads caused a reduction in detected splicing factors, whereas the recovery of MTREX was not affected. These observations all hint at physical properties of NRDE2-associated spliceosomes that make detection difficult in ordinary workflows, such that they have remained unnoticed.

In conclusion, we present evidence for a splicing mechanism that has evolved to effectively select and splice weak 5’SSs in hundreds of genes in mESCs. Investigations of its structural and functional properties, cell type specificity, and evolutionary conservation will further our understanding of pre-mRNA splicing regulation, both in healthy and diseased states.

## ACKNOWLEDGMENTS

We thank the FMI Functional Genomics, FACS, and Microscopy facilities, and Sarah H Carl for support with computational analyses. We thank Philipp Krastel and Mathias Frederiksen (Novartis Institutes for BioMedical Research) for sharing the splicing inhibitor Thailanstatin A. This work was supported by the Swiss National Science Foundation (SNSF), NCCR RNA & Disease (grant 141735), and the Novartis Research Foundation. We thank members of the Bühler lab for discussions and Guy Riddihough (Life Science Editors) for editing the manuscript.

## AUTHOR CONTRIBUTIONS

M.F. conceived the study, designed and performed experiments, analyzed data and wrote the manuscript. M.S. performed bioinformatic analyses. D.H. and V.I. performed and analyzed mass spectrometry. A.C.T. and F.M. helped with bioinformatic analyses. M.B. conceived the study, obtained funding, discussed results and wrote the manuscript.

---

**Figure S1.**
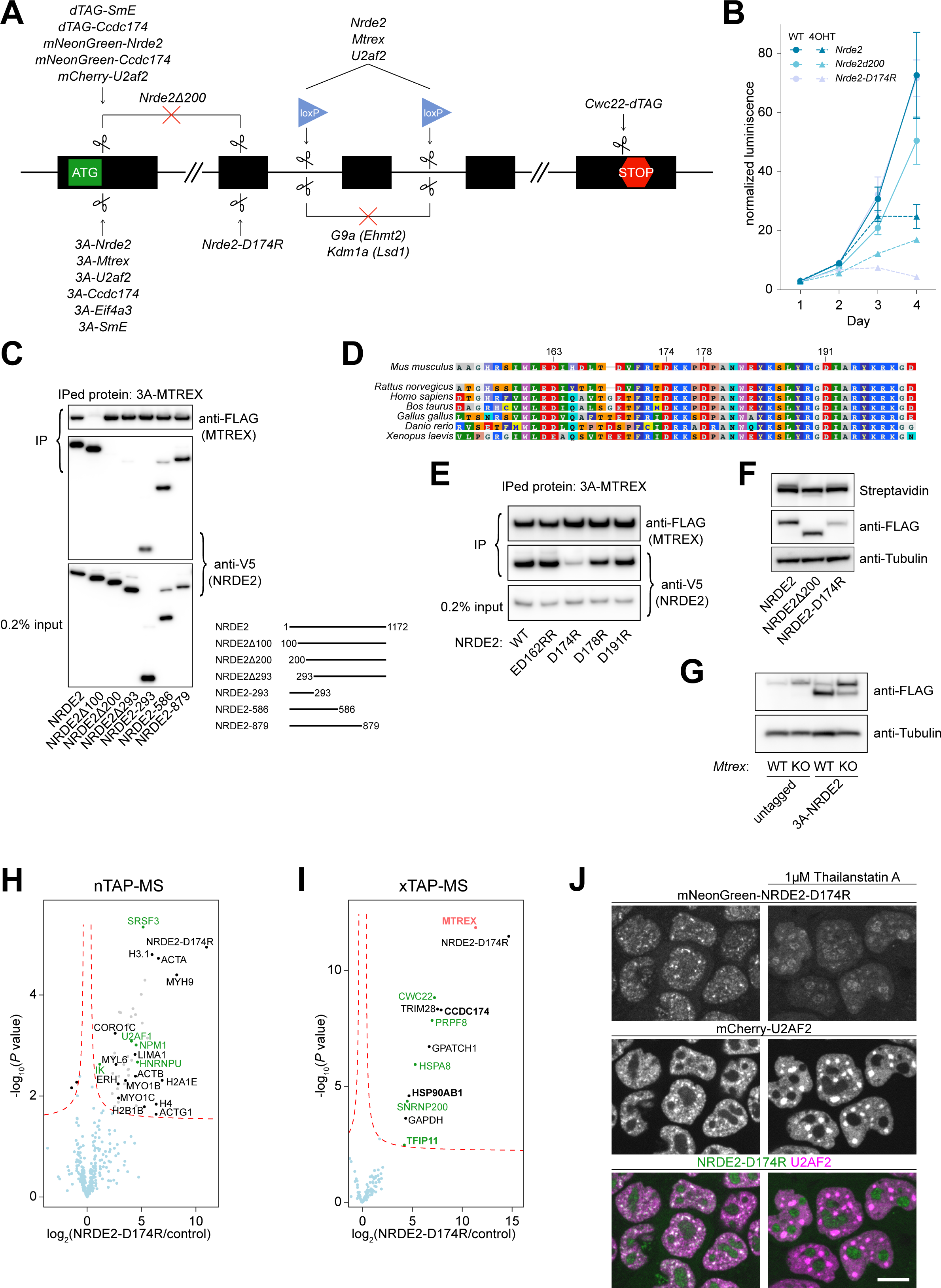
Direct interaction with the MTREX helicase promotes NRDE2 stability, related to Figure 1. (A) A schematic overview of TALEN- or Cas9-mediated genome editing manipulations performed in this study (See also Table S1). (B) Growth curve of 4OHT-inducible knockout *Nrde2*, *Nrde2Δ200* and *Nrde2-D174R* cells grown in the presence or absence of 0.1 μM 4OHT. (C) Streptavidin pulldown and co-immunoprecipitation of endogenous 3A-MTREX from cells transiently overexpressing V5-tagged full length or truncated NRDE2 constructs fused to 2A-Puro. The detection of MTREX by FLAG antibody in NRDE2Δ100 lane was occluded on the reprobed membrane after the initial detection of V5-NRDE2Δ100 protein of identical size. (D) Amino acid sequence alignment of MTREX-interacting region within mouse NRDE2 with its homologs from other vertebrates. (E) Streptavidin pulldown and co-immunoprecipitation of endogenous 3A-MTREX from cells transiently overexpressing V5-tagged wild type or mutant NRDE2 constructs fused to 2A-Puro. (F) Western blotting analysis of endogenous 3A-tagged NRDE2, NRDE2Δ200 and NRDE2-D174R protein expression. The strong band detected in all samples by streptavidin represents an endogenously biotinylated protein. (G) Western blotting analysis of endogenous 3A-tagged NRDE2 expression in conditional *Mtrex*-KO cells untreated (WT) or treated for 3 days with 0.1 μM 4-OHT (KO). The upper band detected by anti-FLAG antibody represents a non-specific target as it is also present in the parental untagged cells. (H) Native TAP-MS analysis of 3A-NRDE2-D174R under low salt conditions (100mM NaCl) compared to untagged control and performed in three independent replicates for each sample. Same labeling order as in Figures 1A and 1B applies. (I) Formaldehyde cross-linking TAP-MS analysis of 3A-NRDE2-D174R compared to untagged control and performed in three independent replicates for each sample. Same labeling order as in Figures 1A and 1B applies. (J) Live cell spinning disk microscopy of endogenous mNeonGreen-tagged NRDE2-D174R with endogenous mCherry-tagged U2AF2. The imaging was performed in untreated cells and cells treated for 6 hours with 1 μM Thailanstatin A. Scale bar 10 μm.

**Figure S2.**
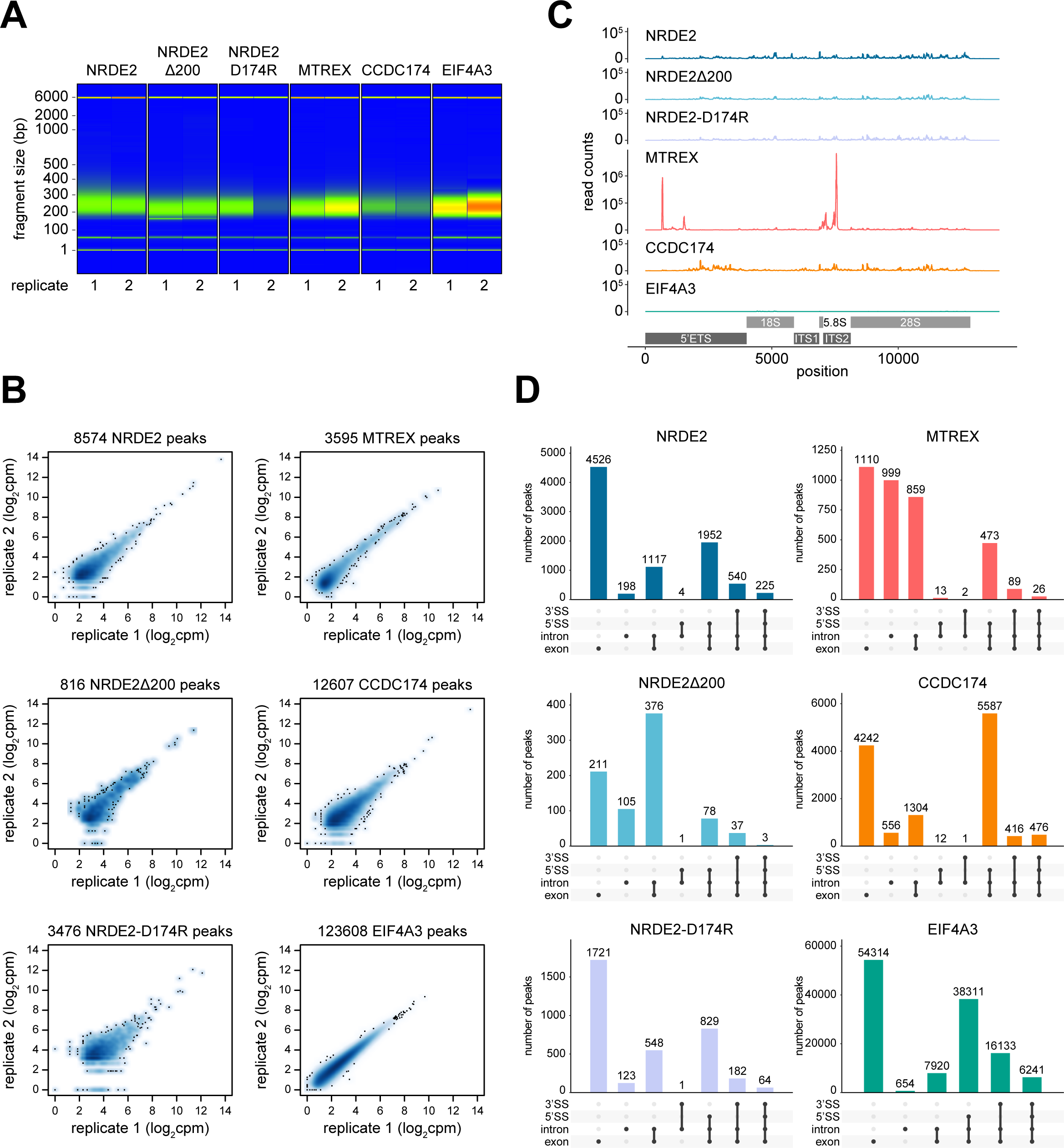
BCLIP-seq allows reproducible detection of low-abundant protein-RNA interactions, related to Figure 2. (A) Gel image from Fragment Analyzer System (Agilent) displaying fragment size distribution of purified BCLIP-seq libraries. The combined size of adaptor sequences is 153 bp. (B) Scatter plots showing the correlation of BCLIP-seq signal between two replicates of each assayed protein. The plotted signal represents normalized read counts in individual peaks identified by a peak calling analysis. (C) BCLIP-seq signal coverage over the first 14 kb of mouse ribosomal DNA repeat unit (GenBank: BK000964). Positions of annotated rRNA segments are provided, ETS = external transcribed spacer, ITS = internal transcribed spacer. (D) UpSet plots displaying the overlap of called BCLIP-seq peaks with different pre-mRNA features and their intersections. Each bar shows a number of peaks overlapping the features marked by a black dot. The features were extracted from the GENCODE transcriptome annotation which may include multiple alternatively spliced transcript isoforms for a single gene. For example, the category exon+intron (third bar) comprises peaks overlapping an alternative exon that is annotated as an exon in some transcript isoforms and as an intron in other isoforms of a given gene.

**Figure S3.**
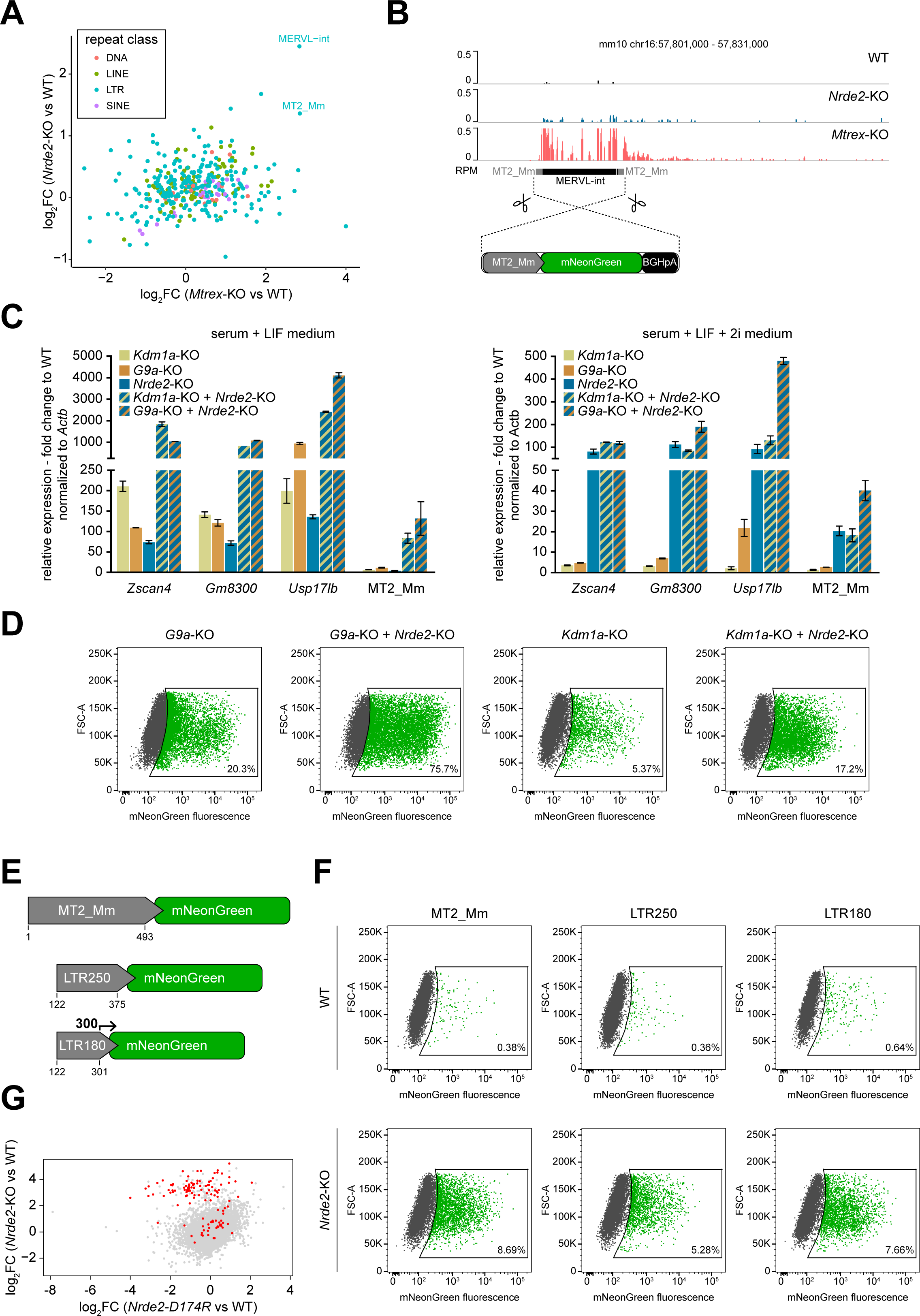
NRDE2 depletion leads to MERVL and 2C gene activation, related to Figure 3. (A) Differential expression of repetitive elements in *Nrde2*-KO and *Mtrex*-KO cells relative to the corresponding WT cells. Each dot represents a unique repeat type based on RepeatMasker annotation. (B) UCSC genome browser screenshot showing normalized RNA-seq tracks at a full length MERVL-containing locus on chromosome 16. Below is a schematic overview of the 2C reporter and its genomic insertion strategy. (C) Quantitative RT-PCR analysis of MT2_Mm and selected 2C marker genes expression in *Kdm1a*-KO, *G9a*-KO, *Nrde2*-KO cells and their combinations relative to WT cells grown in the presence or absence of 2i inhibitors. Plotted is the average of two independent experiments performed in technical triplicates and the error bars represent standard deviation. (D) Flow cytometry analysis of stably integrated 2C reporter expression in *G9a*-KO and *Kdm1a*-KO cells with or without simultaneous induced knockout of *Nrde2* (6 days of 0.1 μM 4OHT treatment). Cells were grown in serum + LIF medium in the absence of 2i inhibitors. (E) Schematic depiction of the 2C reporter variants driven by MT2_Mm fragments of different length. The bent arrow indicates the transcriptional start site determined by 5’RACE. (F) Flow cytometry analysis of the expression of stably integrated 2C reporter variants described in panel (E) in WT and *Nrde2*-KO cells (6 days of 0.1 μM 4OHT treatment). (G) Scatter plot comparing differential gene expression in *Nrde2*-KO and *Nrde2-D174R* cells relative to WT cells. Each dot represents a single gene. Genes upregulated in 2C-like cells (Macfarlan et al., 2012) are highlighted in red.

**Figure S4.**
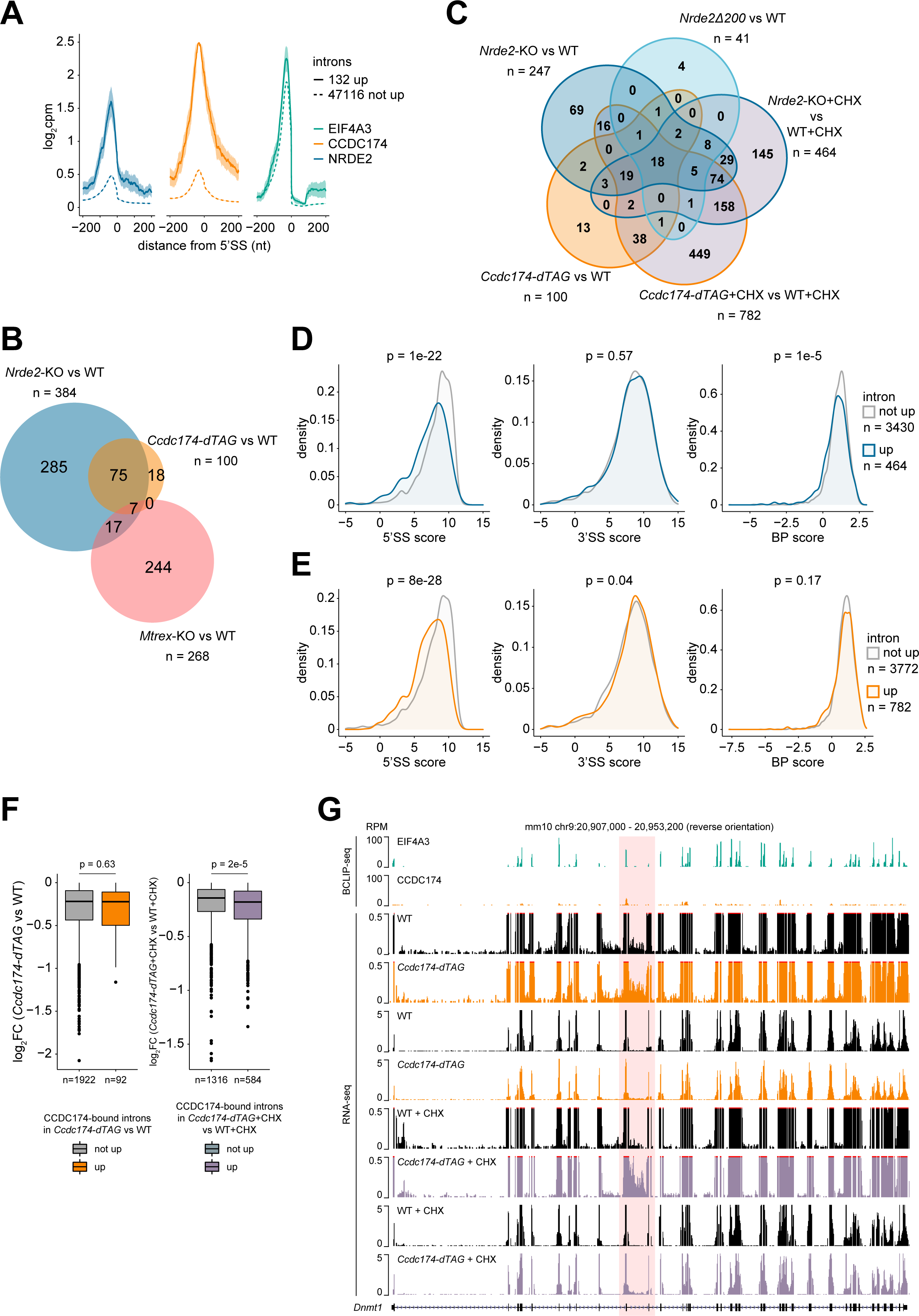
CCDC174 regulates splicing of introns with weak 5’splice sites, related to Figure 3. (A) Metaplots of normalized NRDE2, CCDC174 and EIF4A3 BCLIP-seq signal around 5’SSs of introns upregulated (solid line) or not upregulated (dashed line) in *Ccdc174-dTAG* cells compared to WT. (B) Overlap of CCDC174 target introns (introns with a CCDC174 BCLIP-seq peak at their 5’SS) upregulated in *Nrde2*-KO, *Ccdc174-dTAG* or *Mtrex*-KO cells. (C) Overlap of NRDE2 target introns upregulated in *Nrde2*-KO cells untreated or treated for 4 hours with 100 μg/ml cycloheximide (CHX) and in *Nrde2Δ200* cells with CCDC174 target introns upregulated in *Ccdc174-dTAG* cells untreated or treated with CHX relative to their respective CHX-treated or untreated WT controls. (D) Density plots of the distribution of NRDE2 target introns upregulated (blue line) or not upregulated (grey line) in *Nrde2*-KO cells treated with CHX based on their 5’SS, 3’SS and branch point strength. P-values were calculated using Wilcoxon rank-sum test. (E) Density plots of the distribution of CCDC174 target introns upregulated (orange line) or not upregulated (grey line) in *Ccdc174-dTAG* cells treated with CHX based on their 5’SS, 3’SS and branch point strength. P-values were calculated using Wilcoxon rank-sum test. (F) Differential expression of genes containing CCDC174 target introns in *Ccdc174-dTAG* cells (24 hours of 0.5 μM dTAG-13 treatment) untreated (left) or treated (right) with CHX. Genes are split into two groups: those containing only target introns not upregulated (grey) and those containing at least one target intron upregulated in *Ccdc174-dTAG* cells untreated (orange) or treated with CHX (purple). P-values were calculated using Wilcoxon rank-sum test. (G) UCSC genome browser screenshot showing normalized BCLIP-seq and RNA-seq tracks over *Dnmt1* gene locus. Position of the CCDC174 target intron 10 is highlighted in red.

**Figure S5.**
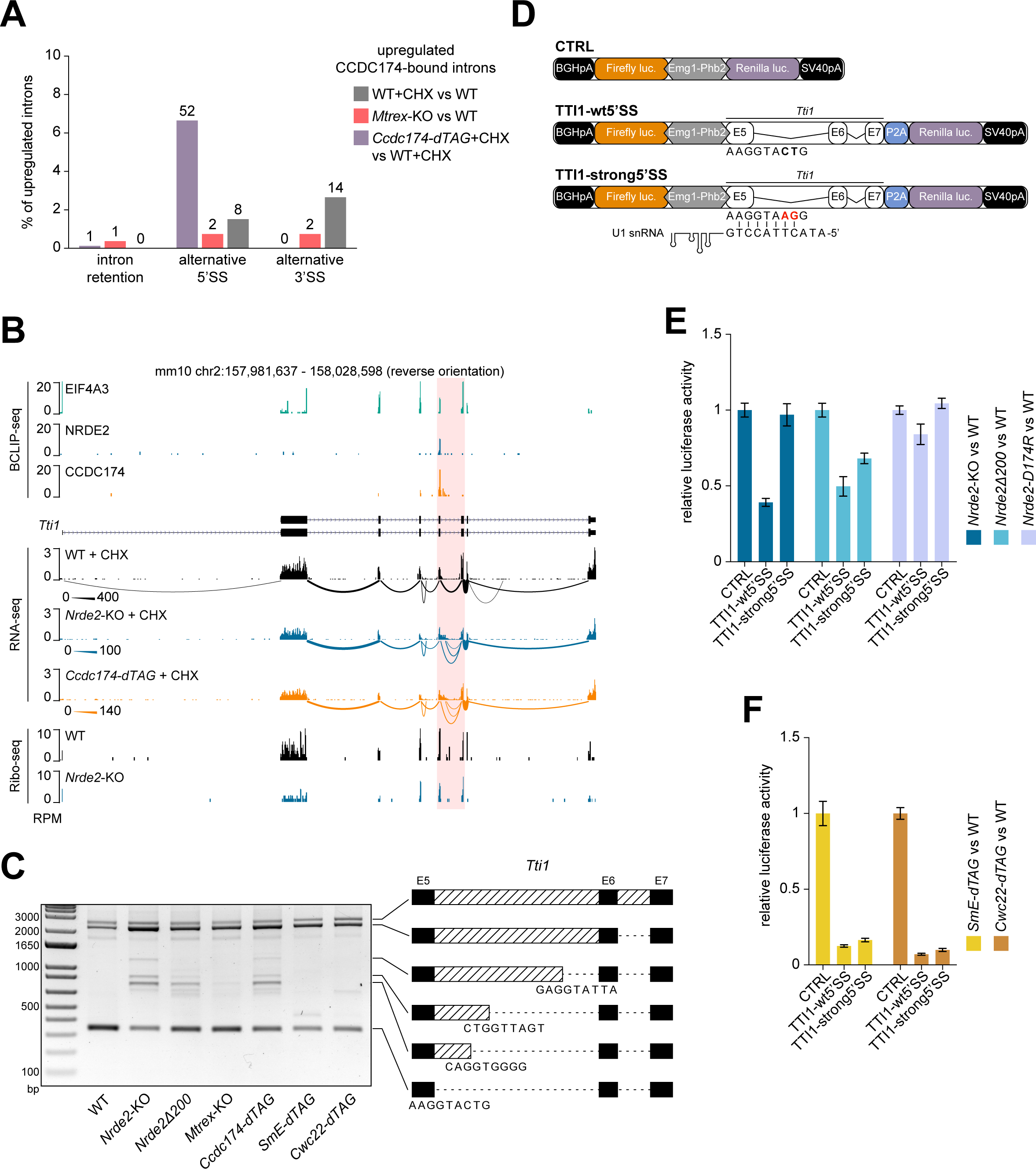
NRDE2 regulates splicing of its target gene Tti1, related to Figure 4. (A) rMATS analysis of alternative splicing events within CCDC174 target introns upregulated in *Mtrex*-KO cells and in CHX-treated WT and *Ccdc174-dTAG* cells relative to their corresponding WT and untreated controls. (B) UCSC genome browser screenshot showing normalized BCLIP-seq, RNA-seq and Ribo-seq tracks over *Tti1* gene locus combined with a Sashimi plot of *Tti1* splicing patterns detected in the RNA-seq data. The thickness of the splice junction connecting lines reflects the number of corresponding spliced RNA-seq reads with a scale provided on the left. NRDE2 and CCDC174 target intron 5 is highlighted in red. (C) RT-PCR analysis of *Tti1* intron 5 splicing with primers binding in exon 5 and 7. The additional bands in *Nrde2*-KO sample were sequenced and the identified cryptic 5’SSs are schematically depicted. (D) Schematic overview of the TTI1 dual luciferase reporters. The mutations introduced in *Tti1* intron 5 5’SS to improve U1 snRNA complementarity in the TTI1-strong5’SS reporter are depicted in red. (E and F) Dual luciferase assay of the transiently transfected TTI1 reporters. Plotted is the average value of normalized Renilla luciferase activity from three independent experiments each performed in triplicate transfections. Error bars represent standard deviation.

**Figure S6.**
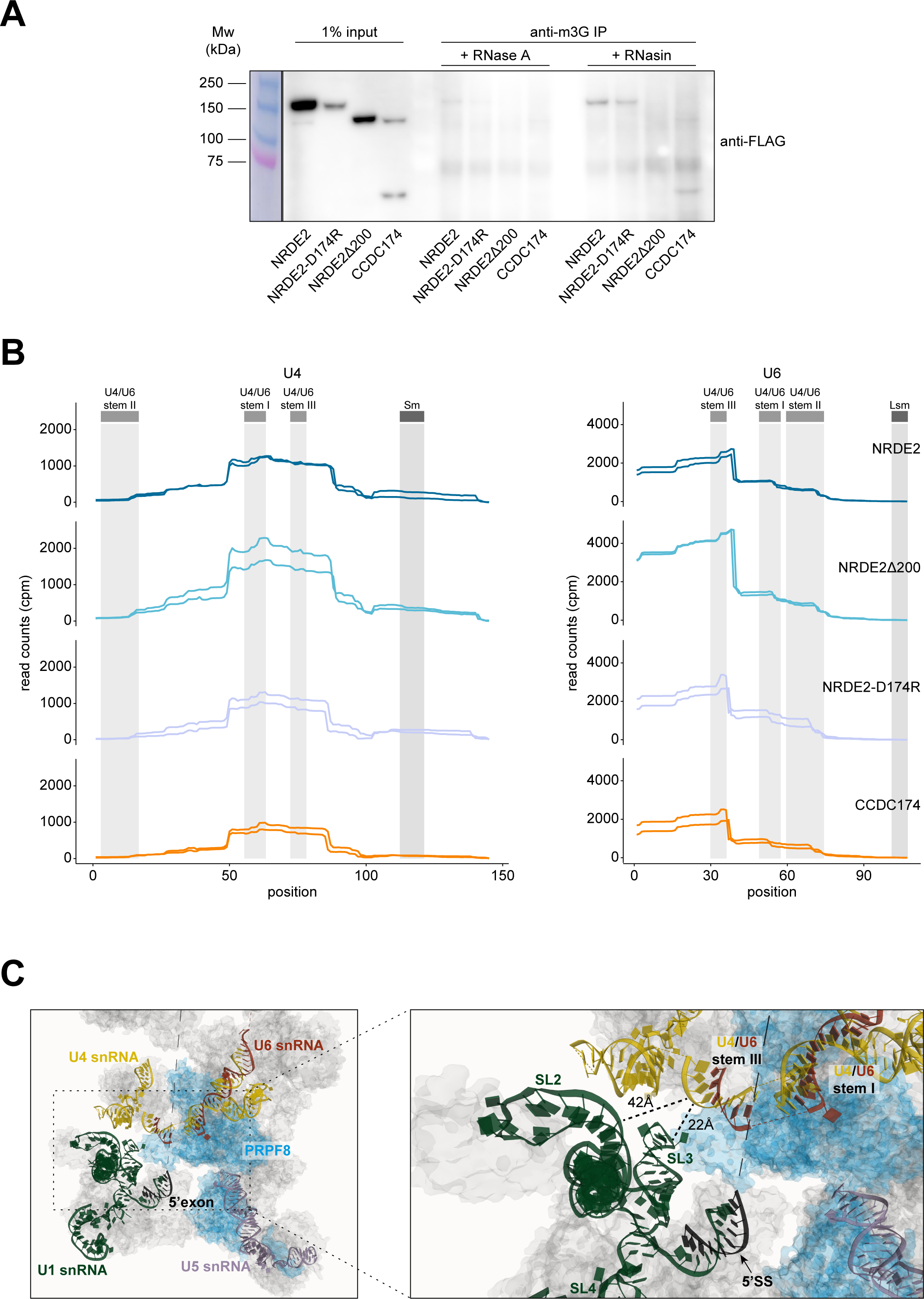
NRDE2 and CCDC174 contacts with snRNAs fit the pre-catalytic spliceosome model, related to Figure 6. (A) Anti-m3G cap pulldown and co-immunoprecipitation of endogenously 3A tagged NRDE2, NRDE2-D174R, NRDE2Δ200 and CCDC174 in the presence of RNase A or the RNase inhibitor RNasin. (B) BCLIP-seq signal coverage over U4 and U6 snRNA sequences showing the two BCLIP-seq replicates of each sample separately. Grey columns indicate the positions of the Sm/LSm ring binding sites and the three U4/U6 base-pairing regions. (C) Snapshot of the human spliceosomal pre-B complex structure PDB ID: 6QX9 (Charenton et al., 2019) showing the proximity of U1 snRNA stem-loops II and III with U4/U6 snRNA stem III. NRDE2 and CCDC174 xTAP-MS interactor PRPF8 is depicted in blue. The black dotted lines in the close-up image on the right depict the closest distance between U1 stem-loops II and III and U4/U6 stem III with the measurement value included.

## METHODS

### Experimental Model and Subject Details

Male mouse embryonic stem cells of a mixed 129 x C57BL/6 background with a heterozygous integration of only Cre-ERT2 (cMB052) or both Cre-ERT2 and V5-tagged BirA (cMB063) in the *Rosa26* locus (Flemr and Bühler, 2015; Ostapcuk et al., 2018) were used as parental lines to generate all the genome-edited cell lines used in this study.

### Cell culture and treatment

Unless stated otherwise, cells were grown on gelatin-coated dishes in S-L-2i medium (DMEM (Gibco) supplemented with 15% fetal bovine serum (Gibco), 1 × non-essential amino acids (Gibco), 1 mM sodium pyruvate (Gibco), 2 mM L-glutamine (Gibco), 0.1 mM 2-mercaptoethanol (Sigma), 50 μg/ml penicillin, 80 μg/ml streptomycin, 1:500 MycoZap Prophylactic (Lonza), home-made LIF conditioned medium and 2i inhibitors: 3 μM GSK-3 inhibitor XVI (Sigma) and 10 μM MEK inhibitor PD0325901 (Tocris)). To induce conditional knockout of *Nrde2* and *Mtrex*, 4OHT (Sigma, 1:5000 dilution of 0.5 mM stock dissolved in ethanol) was added to the medium and replaced daily for a total of 4 and 3 days, respectively. Depletion of endogenous proteins fused to the 2xHA-FKBP12^F36V^ domain (dTAG) was achieved by treating the cells with 0.5 μM dTAG-13 compound (Tocris, 1:1000 dilution of 0.5 mM stock solution in DMSO) for 24 hours. For chemical inhibition of splicing, cells were treated with 1 μM Thailanstatin A (provided by P. Krastel and M. Frederiksen from Novartis Institutes for BioMedical Research, 1:1000 dilution of 1 mM stock solution in DMSO) for 6 hours. To block transcription, cells were treated with 5 μg/ml Actinomycin D (Sigma, 1:1000 dilution of 5 mg/ml stock solution in DMSO) for 2 hours. For translation inhibition, 100 μg/ml Cycloheximide (Sigma, 1:1000 dilution of 100 mg/ml stock solution in DMSO) was added for 4 hours.

### Genome editing

We used both TALEN and Cas9 nucleases to generate cell lines with endogenous gene modifications. Endogenous *Nrde2* truncation and point mutation, tagging with 3A tag and loxP site insertions were achieved using single-stranded oligodeoxynucleotides with short homology arms synthesized by Integrated DNA Technologies. Longer inserts flanked by longer homology arms were cloned into pBluescript II KS-plasmid (Agilent Technologies). All TALEN and Cas9 target sequences and donor sequences for homologous recombination are listed in Table S1. Cells grown in medium without 2i inhibitors were seeded 5 x 10^5^ per well of a 6-well plate and transfected with a mixture of 500 ng donor DNA and 500 ng Cas9-2A-Puro/TALEN mix with pRR-Puro reporter (Flemr and Bühler, 2015) in Opti-MEM medium (Gibco) using Lipofectamine 3000 (ThermoFisher Scientific) according to manufacturer’s instructions. The transfected cells were selected by a 24-hour treatment with 2 μg/ml puromycin (Sigma), expanded at a clonal density and the clones were picked and genotyped by PCR. Correct homozygous insertions were verified by Sanger sequencing.

### Growth curve

Cell growth was monitored using the ViaLight Plus cell proliferation and cytotoxicity bioassay kit (Lonza). Triplicates of 500 cells in 0.5 ml medium were seeded on a 24-well plate. After 24 hours, medium was removed from the wells of Day 0 time point and replaced with 50 μl PBS. Next, 10 μl of ViaLight cell lysis reagent was added, mixed by pipetting and the plate was incubated on a rocking platform for 10 minutes at room temperature. 10 μl of the lysate was mixed with 10 μl of AMR reagent in a 96-well solid bottom white assay microplate (Corning) and after 2 minutes incubation at room temperature the luminiscence was measured in Mithras LB 940 microplate reader (Berthold Technologies). In the wells of later time points, the medium was replaced 24 hours after seeding and every following 24 hours with fresh medium with or without 0.1 μM 4OHT. All the later time points were processed for luminiscence measurement as described above.

### nTAP-MS of 3A-tagged proteins

Each nTAP-MS was performed in three independent replicates. Three near-confluent 10 cm dishes (approx. 5 x 10^7^ cells) were used per each replicate sample. The dishes were placed on ice, cells were rinsed with cold PBS and collected by scraping and 2 minutes spin at 3000 x g and 4°C. The pellets were washed with 0.5 % BSA in PBS and then resuspended in 0.5 ml ice-cold B100 buffer (10 mM Tris-HCl pH 7.5, 2 mM MgCl_2_, 100 mM NaCl, 0.5 % Triton X-100) supplemented with 1 x Halt protease inhibitor cocktail (HALT) (ThermoFisher Scientific) and 100 U Benzonase (Sigma). After 30 minutes shaking (500 RPM) at 12°C, lysates were diluted with 450 μl B100 + HALT and centrifuged for 5 minutes at 16000 x g and 4°C.

For each sample, Protein G Dynabeads (ThermoFisher Scientific) equivalent to 20 μl of the original slurry were resuspended in PBS with 0.02 % Tween-20 and coupled with 2 μl of Anti-FLAG M2 antibody (Sigma) rotating for 30 minutes at room temperature. Coupled beads were washed once with PBS + 0.02 % Tween-20 and twice with B100. The beads were then resuspended in 50 μl of B100 + HALT, mixed with the cleared lysates and rotated for 2 hours at 4°C. Following 3 washes with 1 ml of ice-cold B100, FLAG-tagged proteins were eluted by 15 minutes shaking (500 RPM) in 25 μl of B100 + HALT containing 1 mg/ml 3xFLAG peptide (Sigma) at room temperature. The beads were further rinsed with 425 μl of B100 + HALT and the supernatant was combined with the 25 μl of eluate from previous step.

M-280 Streptavidin Dynabeads (ThermoFisher Scientific) equivalent to 20 μl of the original slurry were pre-washed twice with B100, resuspended in 50 μl of B100 + HALT, mixed with the FLAG beads eluates and rotated for 30 minutes at room temperature. The beads were then washed 3 times with 1 ml of B100, resuspended in 0.2 ml of TN buffer (10 mM Tris-HCl pH 7.5, 100 mM NaCl) and transferred to a new tube. While remaining immobilized on the magnetic stand, the beads were rinsed with an additional 1 ml of TN buffer.

On-bead digestion for mass spectrometry was performed as follows. The washed detergent-free beads were resuspended by vortexing in 5 μl of digestion buffer (8 M urea freshly dissolved in 20 mM HEPES pH 8.5, 5 mM TCEP and 10mM chloroacetamide) and sonicated in Bioruptor Plus (Diagenode) at high energy in 30 cycles 10 s ON/10 s OFF. Next, 1 μl of 0.2 mg/ml LysC protease (Promega) in 50 mM HEPES pH 8.5 was added and proteins were pre-digested for 2 hours rotating at room temperature. The urea concentration was then diluted with 17 μl of 50 mM HEPES pH 8.5 and after adding 1 μl of 0.2 mg/ml trypsin (Promega) in 0.2 mM HCl the digestion continued overnight at 37°C with interval mixing at 2000 RPM for 30 s every 15 minutes. Digested proteins were then subjected to mass spectrometry as described below in ’Mass spectrometry analysis’ section.

### xTAP-MS of 3A-tagged proteins

Each xTAP-MS was performed in three independent replicates. For each replicate sample, cells were harvested by trypsinization, placed on ice, counted on Countess II Automated Cell Counter (ThermoFisher Scientific) and 4.5 x 10^7^ cells were collected by centrifugation for 2 minutes at 300 x g and 4°C. The pellets were resuspended in 1.5 ml of room temperature PBS and cross-linking started by the addition of 1.5 ml of 0.2 % formaldehyde in PBS. After 10 minutes rotating at room temperature, 150 μl of 2.5 M glycine was added and samples were placed on ice for 2 minutes to quench the excess formaldehyde. Cells were pelleted, rinsed with 0.2 % BSA in cold PBS and finally resuspended by vortexing in 180 μl of TMS buffer (10 mM Tris-HCl pH 8.0, 1 mM MgCl_2_, 1 % SDS, 1 x HALT) pre-chilled at 12°C and supplemented with 100 U Benzonase. Following 30 minutes incubation at 12°C, the lysates were diluted with 1.57 ml of DIL. mix containing 4:5 (vol/vol) ratio of H_2_O and dilution buffer (20 mM Tris-HCl pH 8.0, 1 M NaCl, 2 % Triton X-100, 20 mM EDTA) supplemented with 1 x HALT and incubated for 5 minutes on ice. Any remaining insoluble material was removed by a 5 minutes spin at 16000 x g and 4°C.

The cleared lysates were mixed with anti-FLAG antibody-coupled Protein G Dynabeads, which were prepared essentially as described above in the nTAP-MS section, but pre-washed with 2 x 1 ml of WASH buffer (10 mM Tris-HCl pH 8.0, 0.5 M NaCl, 1 % Triton X-100, 0.1 % SDS, 1 mM EDTA) and resuspended in 50 μl of DIL. mix. After 2 hours rotating at 4°C, the beads were washed 3 times with 1 ml of cold WASH buffer and FLAG-tagged proteins were eluted by 15 minutes shaking (500 RPM) in 25 μl of TEDS buffer (10 mM Tris-HCl pH 8.0, 1mM EDTA, 10 mM DTT, 1 % SDS, 1 x HALT) containing 1 mg/ml 3xFLAG peptide at room temperature. The beads were rinsed with another 25 μl of TEDS and the supernatant was combined with the eluate from the previous step and diluted with 0.4 ml of DIL. mix. Next, M-280 Streptavidin Dynabeads (pre-washed 2 x with 1 ml of WASH buffer and resuspended in 50 μl of DIL. mix) were added and the mixture was rotated for 30 minutes at room temperature. The beads were washed 3 times with 1 ml of WASH buffer, resuspended in 100 μl of TEN buffer (10 mM Tris-HCl pH 8.0, 1 mM EDTA, 150 mM NaCl) and transferred to a new tube. The beads immobilized on the magnetic stand were rinsed with an additional 1 ml of TEN buffer. On-bead protein digestion followed exactly as described above in the nTAP-MS section.

### Mass spectrometry analysis

Digested peptides were acidified with 0.8% TFA (final) and analyzed by LC–MS/MS on an EASY-nLC 1000 (Thermo Scientific) with a two-column set-up. The nTAP-MS peptides were applied on an Acclaim PepMap 100 C18 trap column (75 μm ID x 2 cm, 3 μm, Thermo Scientific) in 0.1% formic acid, 2% acetonitrile in H_2_O at a constant pressure of 80 MPa and separated by a linear gradient of 2% – 6% buffer B in buffer A for 3 min, 6% – 22% for 40 min, 22% – 28% for 9 min, 28% – 36% for 8 min, 36% – 80% for 1 min and 80% buffer B in buffer A for 14 min (buffer A: 0.1% formic acid; buffer B: 0.1% formic acid in acetonitrile) on an EASY-Spray column ES801 (50 μm ID x 15 cm, 2 μm, Thermo Scientific) mounted on a DPV ion source (New Objective) connected to an Orbitrap Fusion (Thermo Scientific) at 150 nl/min flow rate. The xTAP-MS peptides were applied on a µPAC trapping column (PharmaFluidics) in 0.1% formic acid, 2% acetonitrile in H2O at a constant flow rate of 5 µl/min and separated by a linear gradient of 3% – 6% buffer B in buffer A for 4 min, 6% – 22% for 55 min, 22% – 40% for 4 min, 40% – 80% for 1 min, and 80% buffer B in buffer A for 10 min on a 50 cm µPAC column (PharmaFluidics) mounted on an EASY-Spray source (Thermo Scientific) connected to an Orbitrap Fusion LUMOS (Thermo Scientific) at 500 nl/min flow rate. Data were acquired using 120,000 resolution for the peptide measurements in the Orbitrap and a top T (3 s) method with HCD fragmentation for each precursor and fragment measurement in the ion trap following the manufacturer guidelines (Thermo Scientific).

Peptides were identified with MaxQuant version 1.5.3.8 using the search engine Andromeda (Cox et al., 2011). The mouse subset of the UniProt version 2019_04 combined with the contaminant DB from MaxQuant was searched and the protein and peptide FDR values were set to 0.05. Statistical analysis was done in Perseus version 1.5.2.6 (Tyanova et al., 2016). Results were filtered to remove reverse hits, contaminants and peptides found in only one sample. Missing values were imputed and potential interactors visualized in the volcano plots were determined using two-sided t-test.

### Co-immunoprecipitation with Streptavidin

Cells were seeded 3 x 10^5^ in 2 ml of S-L-2i medium per well of a 6-well plate and transfected with a mixture of 200 ng of plasmid DNA, 0.6 μl of Lipofectamine 3000 and 0.4 μl of P3000 reagent in 100 μl of Opti-MEM medium. After 24 hours, cells were rinsed twice with ice-cold PBS and lysed directly in the well with 900 μl of CoIP buffer (20 mM Tris-HCl pH 7.5, 150 mM NaCl, 1 mM EDTA, 1 % Triton X-100) supplemented with 1 x HALT and, where indicated, also with 1 μl of 10 mg/ml RNase A (Roche). Following one hour agitation at 4°C, the lysates were collected and centrifuged for 5 minutes at 16000 x g and 4°C. Two μl of the cleared supernatants was transferred to a separate tube as the input sample and the rest was mixed with 100 μl of CoIP buffer containing M-280 Streptavidin Dynabeads. Prior to use, the beads (20 μl of the original slurry) were pre-blocked for one hour at room temperature in PBS containing 0.5 % cold water fish gelatin (Sigma) and 0.02 % Tween-20 and washed twice with CoIP buffer. Following one hour incubation at 4°C, the beads were separated from the lysate and washed four times with 1 ml of CoIP buffer before resuspending in 15 μl of 1 x loading buffer (NuPAGE LDS sample buffer (4 x) and NuPAGE sample reducing agent (10 x) (ThermoFisher Scientific) diluted to 1 x with H_2_O). 13 μl 1x loading buffer was also added to the input samples. All samples were heated for 3 minutes at 95°C and subjected to western blotting as described in the Western blotting section below.

### Co-immunoprecipitation with anti-m3G antibody

Cells were harvested by trypsinization and counted. For each sample, one million cells were rinsed once with PBS and collected by centrifugation at 200 x g for 3 minutes. Cell pellets were resuspended in 1 ml of CoIP buffer supplemented with 1 x HALT and incubated for 10 minutes on ice. The lysates were centrifuged for 5 minutes at 16000 x g and 4°C. Ten μl of the supernatant was transferred to a separate tube as the input sample and stored at 4°C for the duration of the IP. The remaining supernatant was split in two halves, supplemented with 2 μl of anti-2,2,7-trimethylguanosine (anti-m3G) antibody (Sigma) and 1 μl of either RNasin RNase inhibitor (Promega, 40 U/μl) or 10 mg/ml RNase A (Roche) and incubated overnight at 4°C with mixing. The next day 50 μl of Protein G Dynabeads (equivalent to 20 μl of the original slurry, pre-washed 2 x with 1 ml of CoIP buffer) resuspended in CoIP buffer with 1 x HALT were added and incubated for 4 hours at 4°C with mixing. Beads were then washed 3 times with 1 ml of cold CoIP buffer before resuspending in 15 μl of 1 x loading buffer. The inputs were mixed with 4 μl NuPAGE LDS sample buffer (4 x) and 1.6 μl NuPAGE sample reducing agent (10 x). All samples were heated for 3 minutes at 95°C and subjected to western blotting as described in the Western blotting section below.

### Western blotting

Cells were harvested by trypsinization and counted. One million cells were rinsed once with PBS and collected by centrifugation at 200 x g for 3 minutes. Cell pellets were resuspended in 100 μl TMNS buffer (10 mM Tris-HCl pH 8.0, 1 mM MgCl_2_, 100 mM NaCl, 1 % SDS) containing 25 U Benzonase and incubated for 5 minutes at room temperature. 10 μl of the resulting lysate was mixed with 4 μl NuPAGE LDS sample buffer (4 x) and 1.6 μl NuPAGE sample reducing agent (10 x), incubated for 3 minutes at 95°C and loaded on NuPAGE 4 - 12 % Bis-Tris Mini Protein Gel (ThermoFisher Scientific). Proteins were separated by electrophoresis in NuPAGE MOPS SDS running buffer and transferred in Bjerrum Schafer-Nielsen transfer buffer (48 mM Tris, 39 mM glycine, 20 % methanol) onto Immobilon-P PVDF membrane (Millipore) using Trans-Blot SD Semi-Dry Transfer Cell (Bio-Rad) at constant 15 V for 15 minutes, followed by constant 25 V for 25 minutes. We note that for full length NRDE2, using the rapid Trans-Blot Turbo Transfer System (Bio-Rad) resulted in suboptimal transfer that cannot be explained by the protein size. For visualization of biotinylated proteins, the membrane was blocked for 15 minutes at room temperature in TBST containing 1 % BSA before adding HRP-Streptavidin (Sigma) to 1:20000 dilution and incubating for further 30 minutes. The membrane was washed 3 times (5 minutes each) with TBST, rinsed with PBS and the proteins were visualized with Immobilon Western Chemiluminiscent HRP substrate (Merck/Millipore, used at 1:1:3 (H_2_O) dilution) using Amersham Imager 600 (GE Healthcare). For antibody-based western blotting, the membrane was blocked for 10 minutes at room temperature in TBST containing 1 % non-fat dry milk (TBST-NFDM) and then incubated with primary antibody diluted in TBST-NFDM with 0.05 % NaN_3_ at 4°C overnight. The membrane was washed 3 times (5 minutes each) with TBST and incubated for one hour at room temperature with HRP-conjugated secondary antibody diluted 1:10000 in TBST-NFDM. After that, the membrane was washed and the signal was developed as described above. When re-probing the same membrane with a different antibody, the membrane was stripped twice for 5 minutes in 25 mM glycine pH 2.0, with the first incubation supplemented with 1 % SDS, before rinsing thoroughly with water and repeating the blocking and antibody incubations as described above.

### Yeast two-hybrid screen

The yeast two-hybrid screen was performed as a commercial service by Hybrigenics S.A. (Paris, France). A bait cDNA of full length mouse *Nrde2* (encoding amino acids 2-1172) was probed against Mouse Embryonic Stem Cell_RP1 cDNA library with a total of 58.2 million interactions tested. Gene Ontology terms of 87 high confidence interactors (PBS score A - D) were analyzed using the clusterProfiler R package (Yu et al., 2012).

### Live cell imaging

Ibidi μ-Slide 8-well chambers were coated with 10 μg/ml Biolaminin LN-511 (BioLamina) diluted in PBS with 1 mM CaCl_2_ and 0.5 mM MgCl_2_ at 4°C overnight. Coated wells were rinsed with PBS and immediately filled with S-L-2i medium to which 4 - 6 x 10^4^ cells were plated. Cells were imaged 24 hours after seeding on a Nikon Ti2-E Eclipse inverted microscope equipped with Yokogawa CSU W1 spinning disk confocal scanning unit, two back illuminated EMCCD iXon-Ultra-888 (Andor) cameras and CFI Plan Apochromat Lambda 100x/1.45 oil immersion objective (Nikon). Fluorescence was excited with 488 nm iBeam Smart (Toptica) and 561 nm Cobolt Jive (Cobolt) lasers and images (pixel size 0.13 μm) were acquired using VisiView software (Visitron Systems GmbH) with the following settings: 100% laser intensity, 500 ms exposure time, EMCCD GAIN 100 for all mNeonGreen-tagged proteins and 25% laser intensity, 200 ms exposure time, EMCCD GAIN 100 for mCherry-U2AF2. The cells were kept at 37°C and 5% CO_2_ during all treatments and imaging.

### BCLIP-seq of 3A-tagged proteins

For each BCLIP-seq sample, 1.5 x 10^7^ cells were seeded on a 10 cm dish 24 hours before cross-linking. For low abundant proteins, two or four dishes were seeded per sample and processed separately until pooling two at a time during the last FLAG beads wash and, when starting with four dishes, during the last Streptavidin beads wash as described below. Cells were rinsed once on ice and then covered with 5 ml of cold PBS. Open dishes without a lid were placed on an ice-cold aluminium plate in Stratalinker 2400 (Stratagene) and 254 nm UV light was applied for 30 seconds to cross-link the protein-RNA interactions. After cross-linking, PBS was removed, cells were scraped in 2 x 0.9 ml of 0.2 % BSA in cold PBS and pelleted by a 2 minutes spin at 3000 x g and 4°C. Pellets were resuspended by vortexing in 100 μl of TMS buffer (as in xTAP-MS) pre-chilled at 12°C and supplemented with 100 U Benzonase. Following 30 minutes incubation at 12°C, the lysates were diluted with 0.85 ml of DIL. mix (as in xTAP-MS) and incubated for 5 minutes on ice. Any remaining insoluble material was removed by a 5 minutes spin at 16000 x g and 4°C.

The cleared lysates were mixed with anti-FLAG antibody-coupled Protein G Dynabeads resuspended in 50 μl of DIL. mix (as in xTAP-MS) and incubated for 2 hours rotating at 4°C, followed by washes, 3xFLAG peptide elution and incubation with M-280 Streptavidin Dynabeads exactly as described above for xTAP-MS. The Streptavidin beads were washed 3 times with 1 ml of WASH buffer, once with 0.2 ml of 1 x PAP buffer (50 mM Tris-HCl pH 8.0, 250 mM NaCl, 10 mM MgCl2, 0.02 % Tween-20) and transferred in 0.1 ml of 1 x PAP to a 0.2 ml PCR tube.

The purified cross-linked RNA fragments were polyadenylated by resuspending the beads in 20 μl of 1 x PAP buffer containing 0.1 mM ATP and 2.5 U *E. coli* Poly(A) polymerase (NEB), and incubating for 10 minutes at 37°C with interval mixing (2000 RPM for 15 seconds every 3 minutes). The reaction was stopped by adding 180 μl of TES buffer (10 mM Tris-HCl pH 8.0, 1mM EDTA, 2 % SDS) and the beads were washed two more times with 0.2 ml of TES before resuspending in 20 μl of TET buffer (10 mM Tris-HCl pH 8.0, 1mM EDTA, 0.5 % Triton X-100) with 0.5 μl of 20 mg/ml Proteinase K (Roche). After 30 minutes of protein digestion at 50°C with interval mixing, Proteinase K was inactivated for 3 minutes at 85°C. The beads were separated on a magnet, the supernatant was transferred to a new tube and mixed with 20 μl of home-made SPRI beads (equivalent of 1 ml of Sera-Mag Magnetic SpeedBeads, carboxylated, 1 μm, 3 EDAC/PA5 (GE Healthcare Life Sciences) in 50 ml of binding buffer containing 10 mM Tris-HCl pH 8.0, 1mM EDTA, 2.5 M NaCl, 20 % PEG8000, 0.05 % Tween-20 and 0.05 % NaN_3_) and 40 μl of isopropanol. RNA was precipitated rotating for 10 minutes at room temperature, beads were separated and rinsed on a magnet twice with 0.2 ml 80 % ethanol, dried for one minute and finally RNA was eluted in 15.25 μl of 0.02 % Tween-20.

The purified RNA was mixed with 1 μl of 0.2 μM RA3dT18V primer and 1 μl of 10 mM dNTPs (10 mM each dATP, dCTP, dGTP, dTTP), denatured for 3 minutes at 65°C and quickly cooled on ice. Reverse transcription started by mixing with a master mix consisting of 2 μl of 10 x RT buffer (500 mM Tris-HCl pH 8.3, 750 mM KCl, 30 mM MgCl_2_, 20 mM TCEP), 0.5 μl of RNase Block (Agilent) and 0.25 μl of SuperScript II (ThermoFisher Scientific) and incubating at 42°C for 15 minutes, after which 1 μl of 40 mM MnCl2 and 1μl of 20 μM template-switching oligo (TSO) were added and the 42°C incubation continued for another 30 minutes, followed by a 15 minute heat inactivation at 70°C. The reaction was slowly cooled down to room temperature with a Δ -1°C / second gradient and 0.5 μl of each RNase H (NEB, 5 U/μl), RNase T1 (ThermoFisher Scientific, 1000 U/μl) and Exonuclease I (NEB, 20 U/μl) were added to degrade the excess TSO and RT primer for 20 minutes at 37°C, followed by a 15 minute heat inactivation at 80°C. The resulting cDNA was purified by isopropanol precipitation on SPRI beads as described above and eluted in 17.2 μl of 0.02 % Tween-20.

To amplify the cDNA library, the eluates from previous step were transferred to a 0.2 ml low-profile PCR tube (Bio-Rad) and mixed with a master mix consisting of 20 μl of 2 x NEBNext Ultra II Q5 Master Mix (NEB), 1 μl of 40 μM P5* primer, 1 μl of 40 μM PE* primer and 0.8 μl of 100 μM EvaGreen Fluorescent DNA Stain (Jena Bioscience). The reactions were amplified in a CFX96 Real-Time PCR instrument (Bio-Rad) with an initial denaturation at 98°C for 45 seconds, followed by cycles of 98°C for 10 seconds, 65°C for 50 seconds, plate read and 65°C for 10 seconds. The amplification was monitored in real time and the reactions were removed after three cycles of exponential fluorescence signal increase (typically between 12 - 18 total cycles). The amplified library was purified by mixing with 32 μl of home-made SPRI beads and incubating for 5 minutes at room temperature. The beads were then separated and rinsed on a magnet twice with 0.2 ml 80 % ethanol, dried for one minute and DNA was eluted in 20 μl of H_2_O. Equal amounts of libraries generated with different barcode TSOs were pooled together and sequenced on the Illumina HiSeq2500 platform (50 nt single-end reads).

All BCLIP-seq-related oligonucleotides were ordered as standard desalted RNA/DNA oligos or ultramers from IDT and their sequences are listed in Table S3.

### RNA-seq

Total RNA was isolated from near-confluent 6 cm dishes using the Absolutely RNA Miniprep kit (Agilent) including an on-column DNase treatment step according to the manufacturer’s instructions. Libraries were prepared using the TruSeq Stranded Total RNA Library Prep Human/Mouse/Rat kit (Illumina) including a ribosomal RNA depletion step and sequenced on the Illumina HiSeq2500 platform (50 nt single-end reads).

### Quantitative RT-PCR

Total RNA was isolated as described in the RNA-seq section and 500 ng was reverse transcribed using the PrimeScript RT Master Mix (Takara). A volume of cDNA corresponding to 5 ng of the input RNA was subjected to quantitative PCR (qPCR) in a 10 μl reaction using the SsoAdvanced Universal SYBR Green Supermix (Bio-Rad) and 0.4 μM of each forward and reverse primer. The qPCR was performed in a CFX96 Real-Time PCR instrument (Bio-Rad) with an initial denaturation at 95°C for 30 seconds, followed by 40 cycles of 95°C for 5 seconds, 60°C for 10 seconds and plate read. The relative RNA levels were determined using the ΔΔCt method and normalized to *Actb* expression. All qPCR primer sequences are listed in Table S4.

### RT-PCR of splicing isoforms

For the splicing analysis of endogenous transcripts, cells were grown to near confluency on 6 cm dishes and total RNA was extracted using the RNAzol RT reagent (Sigma) following the manufacturer’s protocol. One μg of the RNA was reverse transcribed in a 10 μl reaction using SuperScript IV (ThermoFisher Scientific) and random primers (Agilent) and a volume of cDNA corresponding to 5 ng of the input RNA was subjected to 30 cycles of PCR with the 2 x NEBNext Ultra II Q5 Master Mix (NEB).

To determine the splicing pattern of the luciferase reporter transcripts, 2 x 10^5^ wild-type cells or 4 x 10^5^ *Nrde2*-KO cells (2 days of 4OHT treatment) were seeded on a 6-well plate, transfected with 500 ng of the corresponding reporter plasmid DNA using Lipofectamine 3000 and grown for 48 hours. Cells were harvested in 0.5 ml of RNAzol RT reagent, supplemented with 1 μl of Polyacryl carrier (MRC) and total RNA was extracted following the manufacturer’s protocol. 500 ng of the RNA was treated with ezDNase (ThermoFisher Scientific) and reverse transcribed using SuperScript IV and random primers. A volume of cDNA corresponding to 2.5 ng of the input RNA was subjected to 35 cycles of PCR with the 2 x NEBNext Ultra II Q5 Master Mix (NEB).

The PCR reactions were purified using home-made SPRI beads (as described in the BCLIP-seq section), resolved on a 1 % agarose gel containing Midori Green Advance stain (Nippon genetics) and visualized using the FastGene FAS-DIGI PRO gel imaging system (Nippon genetics). The quantification of individual band intensities was performed using Fiji (Schindelin et al., 2012)The sequences of primers used for splicing pattern analysis are listed in Table S4.

### Ribo-seq

For each Ribo-seq sample, 2 x 10^6^ cells (3 days of 4OHT treatment for *Nrde2*-KO cells) were seeded on a 6 cm dish and grown overnight. Cells were rinsed and scraped in 2 x 0.5 ml of cold PBS on ice and pelleted by a 2 minute spin at 3300 x g and 4°C. The pellet was resuspended by pipetting in 0.2 ml of Ribo-seq buffer (25 mM HEPES-KOH pH 7.5, 200 mM KOAc, 2 mM MgCl_2_, 1 % NP-40) and incubated for 5 minutes on ice, followed by a 5 minute spin at 7600 x g and 4°C. 100 μl of the supernatant was transferred to a new tube, mixed with 100 μl of H2O and 0.4 μl of 250 U/μl Benzonase and incubated for 30 minutes at 37°C with constant shaking. The nuclease digestion was stopped by adding 0.5 ml of RNAzol RT reagent with 1 μl of Polyacryl carrier and RNA fragments were purified following the manufacturer’s total RNA isolation protocol. The precipitated RNA was resuspended in 5 μl of H_2_O, mixed with 5 μl of 2 x Novex TBE-Urea sample buffer (ThermoFisher Scientific) and denatured for 3 minutes at 70°C. RNA was resolved on a Novex 15 % TBE-Urea gel (ThermoFisher Scientific) in 1 x TBE buffer for 75 minutes at 180 V side-by-side with 20 ng of a 35 nt long single-stranded DNA oligo 5’-ACCACTCGAGTCAAAACAGAGATGTGTCGAAGATG (IDT). The gel was stained in 20 ml of 1 x TBE buffer containing 2 μl of SYBR Gold stain (ThermoFisher Scientific) for 5 minutes at room temperature and a band migrating just above the 35 nt oligo was cut from the Ribo-seq sample lane.

The gel piece was submerged in 100 μl of 400 mM NaOAc pH 5.2 and frozen at -80°C. After thawing the gel-containing tube for 5 minutes at 95°C, the gel piece was crushed with a plastic pestle, 300 μl of 400 mM NaOAc pH 5.2 was added and RNA was extracted from the gel by 3 cycles of 5 minute incubation at 95°C and 20 minute shaking at 22°C. The mixture was cleared by a 5 minute spin through a Corning Costar Spin-X filter (0.45 μm cellulose acetate, Sigma) at 20000 x g and room temperature. RNA was precipitated by adding 1 μl of glycogen (Roche) and 1 ml of ethanol to the flow through and incubating at -20°C for one hour. RNA was pelleted by a 15 minute spin at 16000 x g and 4°C, the pellet was washed twice with 0.5 ml of cold 80% ethanol, dried for 5 minutes at room temperature and finally resuspended in 10 μl of C1 buffer (10 mM Tris-HCl pH 7.5, 1 mM EDTA, 1 M NaCl, 0.02 % Tween-20).

Before the next step, custom Ribo-beads were prepared as follows. Streptavidin MyOne C1 Dynabeads (ThermoFisher Scientific) equivalent to 20 μl of the original slurry were washed twice with 0.2 ml of C1 buffer and resuspended in 20 μl of C1 buffer. Two rRNA depletion oligos were ordered from IDT as 3’-biotinylated standard desalted custom DNA oligos CCGTACGCCACATTTCCCACGCCGCGACGCGC/3BioTEG/ and CAAGACGAACGGCTCTCCGCACCGGACCCCGGTCCC/3BioTEG/, resuspended and mixed to 50 μM each. Before the first use, the mixture was denatured for one minute at 95°C and quickly cooled on ice. 2 μl of the rRNA depletion oligo mix was added to the 20 μl of C1 Dynabeads and incubated for 15 minutes at room temperature with occasional mixing. The beads were washed twice with 0.2 ml of C1 buffer and resuspended in 20 μl of C1 buffer.

The purified RNA fragments were mixed with 10 μl of the Ribo-beads, incubated for 3 minutes at 70°C, followed by 10 minutes at 50°C. Beads were separated on a magnet and the supernatant was mixed with the remaining 10 μl of the Ribo-beads. After another round of 3 minutes at 70°C and 10 minutes at 50°C, the supernatant was mixed with 30 μl of home-made SPRI beads (as in BCLIP-seq) and 50 μl of isopropanol and the RNA was precipitated rotating for 10 minutes at room temperature. The beads were separated and rinsed on a magnet twice with 0.2 ml 80 % ethanol, dried for one minute and finally RNA was eluted in 13.5 μl of 0.02 % Tween-20.

The purified rRNA-depleted RNA fragments were polyadenylated by adding 4 μl of 5 x PAP buffer (as in BCLIP-seq), 2 μl of 1 mM ATP and 0.5 μl of 5 U/μl *E. coli* Poly(A) polymerase (NEB) and incubated for 10 minutes at 37°C with interval mixing (2000 RPM for 15 seconds every 3 minutes). The reaction was stopped by the addition of 20 μl of home-made SPRI beads and 40 μl of isopropanol. Subsequent polyadenylated RNA purification, reverse transcription with template switching, cDNA library amplification and sequencing on Illumina HiSeq2500 were performed exactly as described in the BCLIP-seq section.

### Flow cytometry

Cells from a near-confluent 6-well plate were harvested by trypsinization and pelleted by a 2 minute spin at 300 x g. The pellets were washed once and resuspended in cold PBS. The mNeonGreen fluorescence of the endogenously expressed 2C reporter was measured on a BD LSR II flow cytometer (BD Biosciences) using a 488 nm excitation laser and a FITC filter. The acquired data were analyzed using the FlowJo software.

### Luciferase reporter assays

The backbone control luciferase reporter plasmid was created by placing the Renilla and Firefly luciferase coding sequences with SV40 and BGH polyA sites, respectively, under the control of a minimal endogenous bidirectional promoter driving the expression of mouse *Emg1* and *Phb2* genes, into the pBluescript II KS-plasmid (Agilent Technologies). For in-frame fusion with Cdk2 and Tti1 fragments, a P2A self-cleaving peptide sequence was included to separate the Renilla luciferase. Site-directed mutagenesis was performed by PCR with NEBNext High-Fidelity 2x PCR master mix (NEB) using overlapping primers with 15 nt overhangs carrying the desired mutation (IDT).

Cells were seeded 2 - 6 x 10^5^ per well of a 96-well plate and transfected with 10 ng of the corresponding reporter plasmid supplemented with 90 ng of pBluescript II KS-using Lipofectamine 3000. *Nrde2*-KO cells were treated with 4OHT for 2 days prior to transfection. Where appropriate, the cells were treated with the dTAG-13 compound 24 hours post-transfection. After 48 hours post-transfection, cells were washed with PBS and lysed in 25 μl of Passive lysis buffer (Promega) for 20 minutes at room temperature. 10 μl of the lysate was used to measure luciferase activity by sequential mixing with 10 μl of each substrate from the Dual Luciferase Reporter Assay System (Promega). Luminiscence was measured on a Mithras LB 940 microplate reader (Berthold Technologies). All Renilla luminiscence values were first normalized to Firefly and the normalized activity was then calculated relative to the empty control reporter in wild-type cells.

### RNA immunopreciptiation followed by northern blotting

Cells were harvested by trypsinization and counted. For each sample, 5 x 10^7^ cells were collected by centrifugation for 2 minutes at 300 x g and rinsed 1 x with 1 ml of PBS. Cell pellets were resuspended in 1 ml of cold CoIP buffer (20 mM Tris-HCl pH 7.5, 150 mM NaCl, 1 mM EDTA, 1 % Triton X-100) supplemented with 1 x HALT in a 15 ml Bioruptor tube containing one scoop of sonication beads (Diagenode) and sonicated in Bioruptor Pico (Diagenode) in 5 cycles of 30 s ON/30 s OFF. Lysates were cleared by centrifugation for 5 minutes at 16000 x g and 4°C and 1 μl of the supernatant was taken to a separate tube as 0.1% input. The remaining supernatant was mixed with anti-FLAG antibody-coupled Protein G Dynabeads (prepared as described above in the nTAP-MS section, but pre-washed with 2 x 1 ml of CoIP buffer and resuspended in 50 μl of CoIP buffer with 1 x HALT). After 2 hours rotating at 4°C, the beads were washed 3 times with 1 ml of cold CoIP buffer and FLAG-tagged proteins were eluted by two consecutive 15 minutes incubations with shaking (500 RPM) at room temperature - first in 100 μl of CoIP buffer with 1 x HALT containing 0.25 mg/ml 3xFLAG peptide, second in 25 μl of CoIP buffer with 1 x HALT containing 1 mg/ml 3xFLAG peptide. The beads were rinsed with another 325 μl of CoIP buffer and the supernatant was combined with the eluates from the previous step. M-280 Streptavidin dynabeads (pre-washed 2 x with 1 ml of CoIP buffer) resuspended in 50 μl of CoIP buffer with 1 x HALT were then added and the samples were incubated for 30 minutes at room temperature with mixing. Beads were washed on ice 3 x with 1 ml of cold CoIP buffer and resuspended in 0.5 ml RNA extraction buffer (10 mM Tris-HCl pH 8.0, 10 mM EDTA, 1 % SDS, 350 mM NaCl, 7 M urea), which was also added to the inputs. RNA was extracted by adding 0.5 ml of buffered phenol:chloroform:IAA pH 8.0 (Sigma) and shaking for 1 minute, followed by a 5 minutes spin at 20000 x g and room temperature. The upper phase was transferred to a new tube and RNA was precipitated by mixing with an equal volume of isopropanol in the presence of 1 μl of Polyacryl carrier (MRC). After 10 minutes incubation at room temperature, RNA was pelleted by centrifugation for 30 minutes at 16000 x g and 4°C, washed once with 0.5 ml 75% ethanol, air-dried for 5 minutes and resuspended in 5 μl of water. Next, 5 μl of 2 x Novex TBE-Urea Sample Buffer (ThermoFisher Scientific) was added, samples were denatured for 3 minutes at 95°C and immediately cooled on ice. RNA was separated on a 6% Novex TBE-Urea gel (ThermoFisher Scientific) for 70 minutes at 90 V in 1 x TBE buffer pre-heated to 50°C. The gel was briefly rinsed with 1 x TBE and RNA was transferred to positively charged Nylon membrane (Roche) using Fastblot (Biometra) for 30 minutes at a constant current of 200 mA. After the transfer, RNA was cross-linked to the wet membrane for 90 s using Stratalinker 2400. The membrane was air-dried, washed in 2 x SSC buffer containing 0.1 % SDS and pre-hybridized in 10 ml of ULTRAhyb-Oligo buffer (ThermoFisher Scientific) for 90 minutes at 37°C. Twenty picomoles of a U1 snRNA-specific 3’-end biotinylated ssDNA oligo (GTATCTCCCCTGCCAGGTAAGTAT/Bio/ (Ishikawa et al., 2014)) was then added and hybridized overnight at 37°C. The membrane was washed 3 x for 15 minutes with 50 ml of pre-warmed 2 x SSC buffer containing 0.5 % SDS at 37°C and the biotin signal was detected with Chemiluminiscent Nucleic Acid Detection Module kit (ThermoFisher Scientific) following the manufacturer’s instructions. The signal was visualized using Amersham Imager 600 (GE Healthcare).

### Quantification and Statistical Analysis

#### BCLIP-seq data preprocessing and alignment

BCLIP-seq reads were preprocessed as described previously for CRAC method (Tuck et al., 2020). Briefly, adapters and low quality bases were trimmed and duplicate reads were collapsed. Samples were split according to their barcodes and low complexity regions were removed from the 3’ ends of the reads.

The preprocessed reads were aligned (splicing-aware) to the mouse genome (mm10) including the transcriptome annotation of GENCODE release M23 using STAR version 2.7.0a (Dobin et al., 2013) with the parameters:

--outFilterMultimapNmax 20

--outFilterMismatchNoverLmax 0.05.

For BCLIP-seq read categorization (Figure 2B), non-collapsed reads were aligned to the mouse ribosomal DNA repeating unit (GenBank: BK000964) using Bowtie2 version 2.3.5.1 (Langmead and Salzberg, 2012) with –sensitive parameters. Then the unmapped reads were mapped in the same way to Gencode release M23 protein coding transcripts. Finally, the leftover unmapped reads were mapped with the same parameters to the mm10 mouse genome. Mapped reads were counted using SAMtools flagstat version 1.10 (Li et al., 2009).

To analyze snRNA-mapping reads, non-collapsed reads were aligned to snRNA sequences using Bowtie2 version 2.3.5.1 with –sensitive parameters. Reads that were at least 20 bp long and mapped with less than 1 edit distance (NM:i:0 or NM:i:1) were used to calculate coverage across snRNAs.

#### RNA-seq read alignment

RNA-seq reads were aligned (splicing-aware) to the mouse genome (mm10) including the transcriptome annotation of GENCODE release M23 using STAR version 2.7.3a. with the parameters:

--outFilterMultimapNmax 100

--outFilterMismatchNoverLmax 0.05

--outSAMmultNmax 1

### Selection of expressed 5’ splice sites

Genocde release M23 annotation was used for all analysis unless otherwise stated. The number of reads in all genes for all WT samples were counted using featureCounts from the RSubread package (Liao et al., 2019) with the following parameters: allowMultiOverlap=FALSE, minOverlap=1, countMultiMappingReads=FALSE, fraction=FALSE, minMQS=255, strandSpecific=2.

All genes with FPKM > 0 were used as expressed genes. In addition, transcript abundance was quantified using Salmon version 1.2.0 (Patro et al., 2017) with the parameters:

-l A -r --validateMappings –gcBias

and only transcripts with TPM > 0 were kept. Then introns (and splice sites) were extracted from these transcripts using the GenomicFeatures R package (Lawrence et al., 2013) and annotated using the EnsDb.Mmusculus.v79 R package.

Heatmaps were generated using the MiniChip R package (https://github.com/fmi-basel/gbuehler-MiniChip): Coverage was calculated in regions +/- 200bp around all 5’ splice sites, using only uniquely mapping reads to the forward strand, and displayed as the average cpm across two replicates.

### Analysis of 5’ splice site-overlapping BCLIP-seq reads

The regions +/- 100 bp around 5’ splice sites of introns longer than 1 kb were selected. Those with less than 50 BCLIP-seq reads (in total over all samples) were removed. Each BCLIP-seq.bam file was loaded using RSamtools R package and the reads that overlap 5’SSs were selected. Spliced or unspliced reads were counted separately for each junction and each sample .bam file. Then the 5’SSs with spliced and unspliced read counts above 10 were selected, and the fold change between spliced and unspliced read counts was calculated.

### Differential gene expression analysis

The number of correctly stranded uniquely mapping RNA-seq reads was quantified in genes using featureCounts (RSubread package). Cycloheximide (CHX)-treated and untreated samples were analyzed separately using DEseq2 (Love et al., 2014), first filtering out genes that have less than 200 counts across all samples or 80 counts across all CHX samples. Genes with an adjusted p-value < 0.01 and log_2_FC > 1/-1 were counted as up/downregulated.

### Differential repeat expression analysis

Repeat expression was quantified with featureCounts using RepeatMasker tracks for LTR, LINE, SINE and DNA elements, removing repeats that overlap genes, using only uniquely mapping reads. The counts were normalized using the TMMwsl method from the edgeR package (Robinson et al., 2010), taking the total number of uniquely mapped reads as library size. Repeats with the maximum cpm below 0.5 were filtered out and the limma R package (Ritchie et al., 2015) was used to calculate differential expression. The average log_2_FC across all genomic instances of a given repeat element (with at least 5 genomic locations) was then calculated.

### Differential intron expression analysis

The number of correctly stranded uniquely mapping RNA-seq reads was quantified in expressed introns using featureCounts with the parameters: useMetaFeatures=FALSE, allowMultiOverlap=TRUE, minOverlap=10, countMultiMappingReads=FALSE, fraction=FALSE, minMQS=255, strandSpecific=2 CHX-treated and untreated samples were analyzed separately using DEseq2, first filtering out introns that have less than 110 counts across all samples or 40 counts across all CHX samples. Introns with an adjusted p-value < 0.05 and log_2_FC > 0.5/-0.5 were counted as up/downregulated. Finally, upregulated introns within genes that are not upregulated (log_2_FC < 0) were selected for further analysis.

### Intron features and alternative splicing

All intron features (GC content, length, splicing scores) were calculated using Matt (Gohr and Irimia, 2018). Fastq files of replicates within a group were combined and mapped to the genome using HISAT2 version 2.1.0 (Kim et al., 2019), followed by transcriptome reconstruction using StringTie version 2.1.1 (Pertea et al., 2015). All resulting .gtf files were merged and annotated using GffCompare version 0.11.6 (Pertea and Pertea, 2020). All rMATS (Shen et al., 2014) comparisons were done using the resulting .gtf file. Retained introns, novel 5’SSs or 3’SSs were selected based on FDR < 0.01 and inclusion level difference > 0.

### Peak finding

CLIPper tool (Lovci et al., 2013) was used to find peaks in STAR-mapped collapsed low-complexity-stripped BCLIP-seq .bam files. Peaks that overlapped between the two replicates, were longer than 35 bp and contained more than 5 reads (in both replicates combined) in a 50 bp window around the peak center were kept.

### Ribo-seq analysis

Ribo-seq reads were preprocessed as described for BCLIP-seq. Read mapping and differential gene expression analysis for Ribo-seq data was performed as for RNA-seq data. Genes with less than 10 reads across all samples were excluded from differential expression analysis.

### Materials, Data, and Code Availability

All reagents and cell lines generated in this study are available from the corresponding author, Marc Bühler, upon Materials Transfer Agreement. All sequencing data generated in this study has been deposited at NCBI GEO and is available under accession number GSE179744. Mass Spectrometry data has been deposited at ProteomeXchange via the PRIDE database under accession number PXD029392. Original Western and Northern Blot images have been deposited at Mendeley Data and can be accessed at https://data.mendeley.com/datasets/v24ytccs4h/2. Custom scripts for data analysis are available upon request, other tools used are indicated in the respective Methods sections.

### SUPPLEMENTAL INFORMATION

Table S1. A list of mES cell lines generated and used in this study, including the details of all genome editing steps.

Table S2. A complete list of NRDE2 interactors identified in a yeast two-hybrid screen of full length *Nrde2* cDNA against an mES cell library.

Table S3. Description of oligos used for BCLIP-seq library construction.

Table S4. A list of primers used for qPCR and for splicing pattern PCR analysis.

